# Convergent natural selection at both ends of Eurasia during parallel radical lifestyle shifts in the last ten millennia

**DOI:** 10.64898/2026.04.03.716344

**Authors:** Alison R. Barton, Nadin Rohland, Swapan Mallick, Ron Pinhasi, Ali Akbari, David Reich

## Abstract

Ancient DNA-based studies of natural selection have focused on West Eurasia due to the availability of large sample sizes, but rich insights are expected to come from comparative studies that can reveal which patterns are shared and which region-specific. We test around seven million variants for selection in 1,862 ancient East Eurasians (867 with new data) distributed over the last ten millennia. Using a generalized linear mixed model to control for population structure, we identify 40 genome-wide significant signals of selection, which have a particularly strong impact on immune and cardiometabolic traits just as in West Eurasia. East and West Eurasia show highly correlated signals of adaptation both for individual alleles and for complex traits, showing how these geographically separate groups experienced convergent evolution in response to parallel transitions to food producing economies and the accompanying lifestyle changes. An exception is the genetic determinants of light skin color: West Eurasians depigmented in the last 10,000 years, but most skin lightening in East Asians arose prior to the Holocene.

---

Selection scans using ancient DNA time series have provided insights into traits under selection, including discovery of strong selection affecting risk alleles for clinically important phenotypes such as celiac disease and multiple sclerosis (*1–4*), fatty acid metabolism (*5*), and resistance to leprosy and tuberculosis (*6*). However, these studies have largely been carried out in West Eurasia, and to generalize the findings, it is important to carry out similar analyses in other parts of the world (*7*).

A recently developed method for detecting natural selection based on a Generalized Linear Mixed Model (GLMM) (*8*) enables more effective screens for natural selection in ancient DNA by avoiding the need to model explicitly the demography of the studied populations. By taking advantage of the independent experiments of nature that have occurred at different times and places, this method tests for consistent trends in allele frequency change over time that can only be explained by directional selection: sustained rises in allele frequency due to an increase in fitness in carriers.

We study East Eurasia over the past ten millennia. The advent of rice and millet farming at the beginning of this period led to transformations in lifestyle and diet; the introduction of pastoralism contributed to longer distance movement; the rise of state societies and orders of magnitude of population growth created unprecedented conditions; and, with these changes, new pathogen exposures emerged.

To test for natural selection in response to these changes, we follow the protocol developed in previous work in West Eurasian ancient DNA time series, leveraging three innovations to increase power to detect selection: use of the GLMM, intensive data cleaning, and assembly of a much larger dataset to detect subtle but consistent shifts in allele frequency over time and space. We report data for the first time data from 817 individuals for which we performed in-solution enrichment of ancient DNA libraries for more than a million single nucleotide polymorphisms (SNPs). We co-analyze these sequences with those from 50 individuals for whom we increase data quality on previously reported individuals, and merge with 995 individuals with previously reported data as well as 610 modern East Asians (**Tables S1-3**).

To maximize power to detect selection, we included as many individuals with primarily East Asian ancestry as possible, including not just in the core region of East Asia but also Central Asia. With the GLMM, heterogenous sample sets are expected to yield additional power when selection has pushed in a consistent direction over space and time but lose power when selection has fluctuated. We restrict to samples excavated 5 degrees of longitude or more to the east of the eastern geographical limit of the West Eurasian study (*1*), ensuring no sample overlap (**Fig. 1A**). We further filtered based on ancestry: only keeping individuals with more East Asian ancestry based on their position in Principal Component Analysis (**Materials and Methods; Supplementary Text A**). The analyzed samples have an average of ∼67% East Asian ancestry (median: 84%) (**Supplementary Text C**). We find enrichment of signals of selection among variants significant (*p* < 5e-8) in GWAS and for eQTL and sQTL loci from GTEx (*9*, *10*) in populations of European ancestry (**Fig. 1B**). To assess our signals using an independent dataset with ancestry more relevant to our cohort, we analyzed the combined GWAS results from Biobank Japan (BBJ) (*11*) and the Taiwan Precision Medicine Initiative (*12*) (TPMI). At these positions, we observe a plateau of enrichment of ∼8x past our rescaled significance threshold of |X| > 5.45 (**Materials and Methods**). We also observed an increase of the residual Haplotype Frequency Score (HAF) around the significance boundary of |X| > 5.45 (**Fig. 1C**), providing an independent line of evidence for the signals being real.

**Fig. 1.**
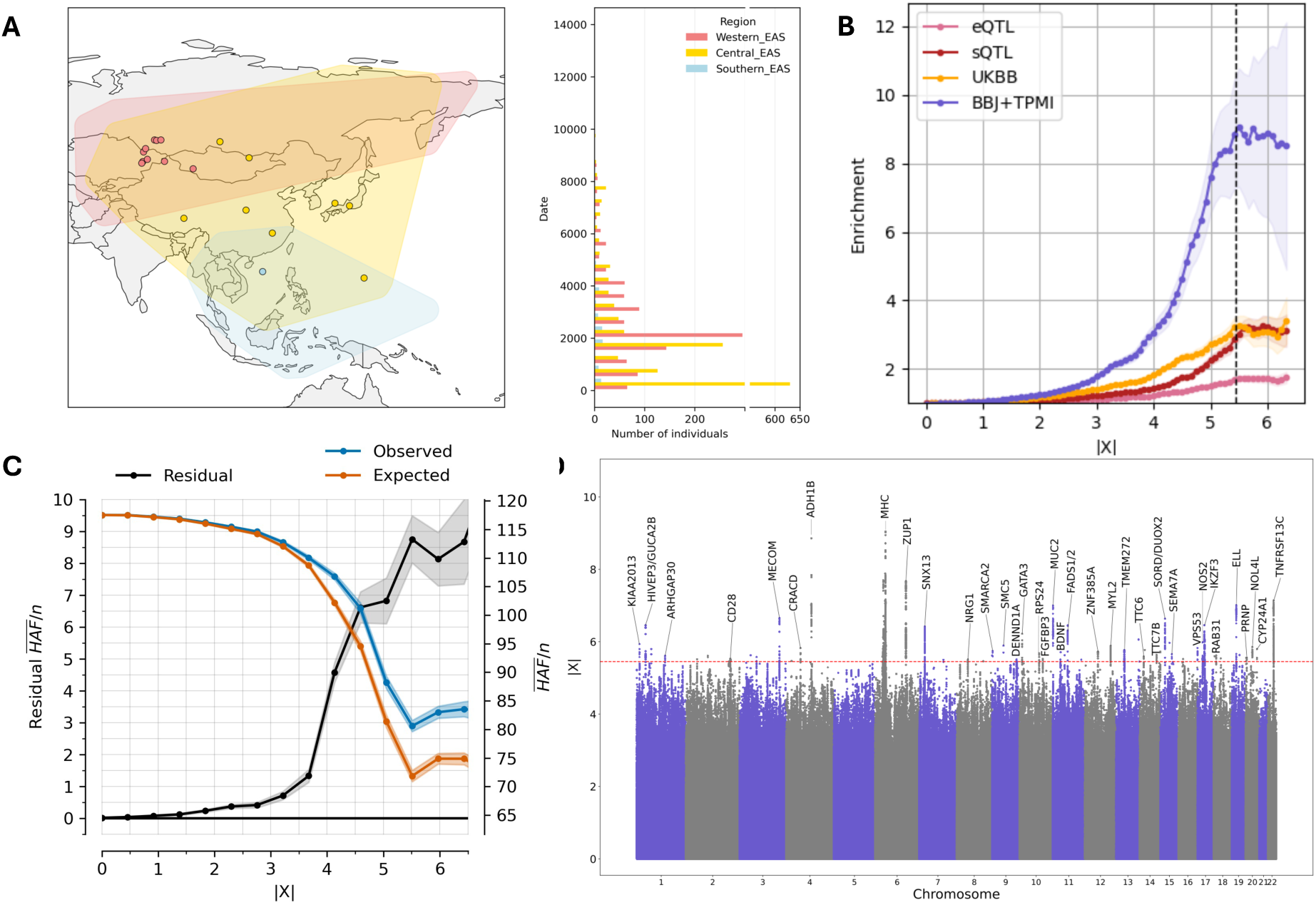
Selection scans using East Eurasian individuals reveal 40 signals of selection that are enriched for biological function. (**A**) Individuals were assigned to a geographic region based on membership in genetic clusters. Points represent the mean location of each genetic cluster included in the analysis. The barplot shows the distribution of samples over time using the same regions. (**B**) Signals of selection are enriched in functional annotations including for GWAS hits from the UK Biobank and from Biobank Japan and the Taiwan Precision Medicine Initiative. (**C**) The residual HAF score also increases with increasing selection statistic. (**D**) Manhattan plot showing the 35 non-MHC independent signals of selection in the East Eurasian cohort.

## Single variants under selection

We identified 35 independent (R^2^ < 0.05), significant signals of selection outside the MHC region and 5 within it (**Fig. 1D**; **Fig. 2; Data S1**), using our |X| > 5.45 threshold of genome-wide significant calibrated based on GWAS data, corresponding to a *p* > 0.99 probability of the alleles being genuinely selected (**Materials and Methods**). Of the non-MHC alleles, 32 have not previously been reported in ancient DNA scans (*7*, *13*), and 25 were not reported in long-range haplotype and allele-frequency tests of modern populations (*14–22*) (**Supplementary Text D**). We highlight a subset of loci of particular interest.

**Figure 2.**
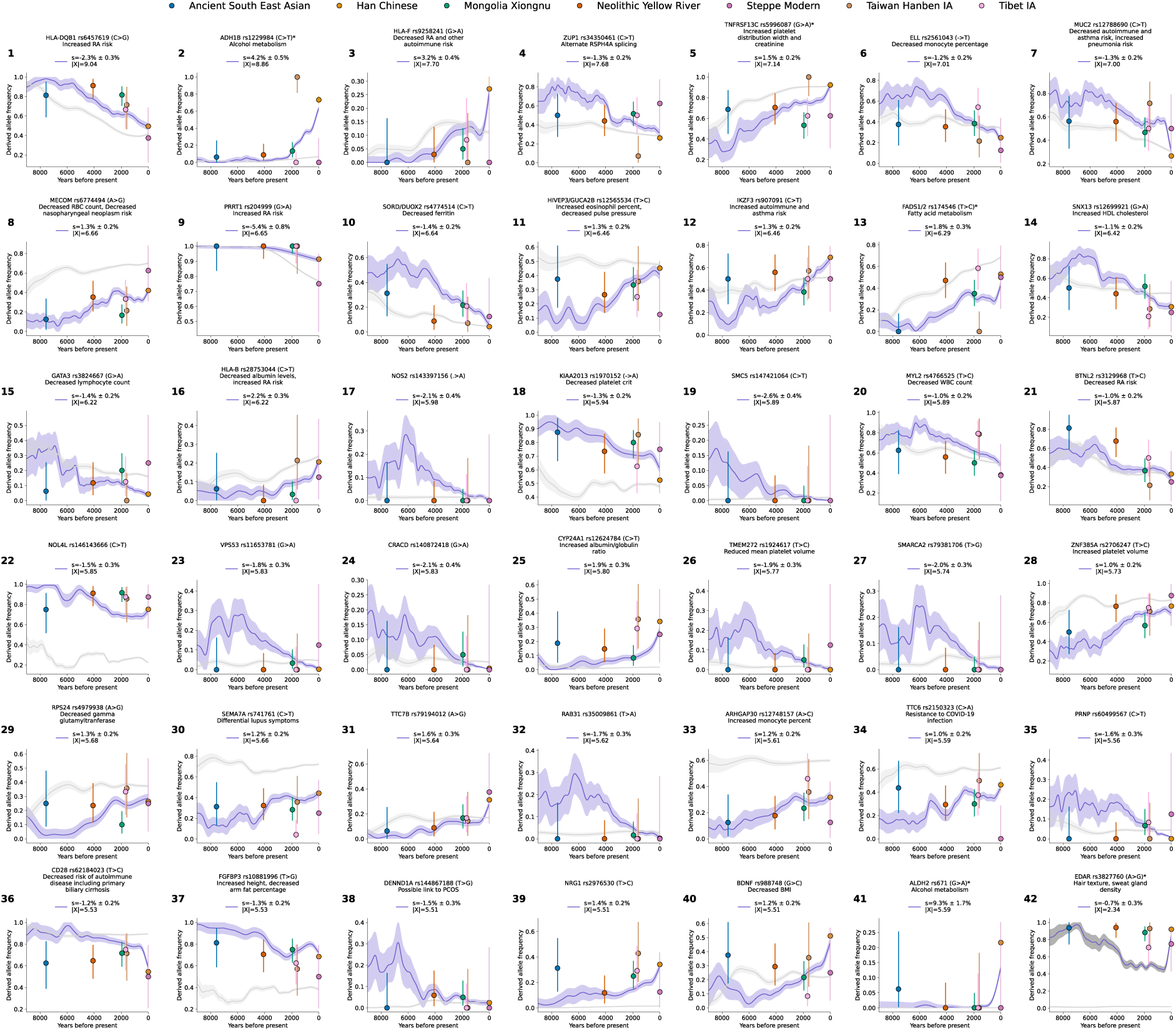
Allele frequency trajectories for the 40 significant, independent signals of selection. The light gray trajectories show the allele frequencies of the same variant in the West Eurasian cohort (*4*) over the same time period. The dots represent the allele frequencies in several relevant populations from the cohort for reference. These samples are included in the scan other than for Ancient South East Asia where several samples not passing QC were included to provide an approximate estimate for this region and time period. **4.41** includes the trajectory for ALDH2 for reference it is R^2^ = 0.13 and D′ = 0.89 with the variant in **4.20** at MYL2. 4.42 shows the trajectory for EDAR, which commonly appears in selection scans of East Asia. We observe it is already at high frequencies at 8,000 years ago and any variation in frequency after this point in our cohort is explainable by population structure. We use the following abbreviations: IA, iron age; RA, rheumatoid arthritis; PCOS, polycystic ovarian syndrome; WBC, white blood cell count; BMI, body mass index.

### New insights for alleles with previous reports of selection in East Asia

#### ADH1B

The strongest signal outside the HLA region is a missense variant related to alcohol metabolism (rs1229984 C>T) previously reported as under selection in East Eurasia (*14*, *23–27*) (**Fig. 2.2**). The *ADH1B* gene encodes an alcohol dehydrogenase protein responsible for converting ethanol into acetaldehyde. The variant common in East Asia and under positive selection here increases the efficiency of this step leading to more rapid accumulation of acetaldehyde (*28*). The allele frequency trajectory shows an abrupt rise in frequency roughly 4000-3000 years before present (BP) with a selection coefficient of *s* = 4.18% ± 0.39% (X = 8.86). Rice fermentation occurred in the archaeological record much earlier (10000-9000 BP) (*29*), suggesting a relatively late onset of selection, supporting theories (*30*) of a rise beginning around 2,800 years ago rather than models of a rise at the same time as the mutation appeared (*31*). In West Eurasia, we also detect significant selection over the same period (∼3000-2000 BP), albeit weaker (*s* = 2.31% ± 0.11%; X = −8.82) and reaching only MAF < 5% in modern populations. This particular variant is dropped from the West Eurasian analysis as it was flagged as low-quality in that cohort; however, a tag SNP (R^2^ = 0.54; D′ = 1 in 1000 Genomes Project European ancestry samples) for the causal missense variant, rs145452708, is one of the 479 hits in ref. (*1*) with 4.33 ± 0.53% and X = 8.14 (*32*, *33*); previous studies also demonstrated selection in East Africa on the variant from West Eurasia further supporting convergent evolution (*34*).

#### MYL2 and ALDH2

We also observe selection at a second variant related to alcohol metabolism (**Fig. 2.20,2.41)**. The lead variant, rs4766525 (T>C), is in weak linkage disequilibrium (R^2^ = 0.13; D′ = 0.89) with rs671 (G>A) in our dataset, the causal variant in *ALDH2* for altered alcohol metabolism today. Selection at the broader locus appears as early as 6000 years ago (s = −1.04% ± 0.15%; X = −5.89, but rs671 does not appear at appreciable frequency until ∼2000 years ago, with most of the rise occurring during the last 1000 years (*s* = 9.31% ± 1.67%; X = 5.59). rs4766525 is also under selection in West Eurasia (*s* = −0.80% ± 0.09%.; X= −8.83) and is associated with decreased white blood cell counts in UK Biobank data, likely related to the nearby gene *SH2B3*. Given the weak LD and distinct timings, these two variants likely tag two separate selection processes at this locus. Taken together with *ADH1B*, these findings support a relatively recent onset of selective pressures that led to changes in alcohol metabolism, shared in part with West Eurasia. We note that in our Taiwan Hanben Iron Age reference set, *ADH1B* is at apparent fixation in the population (mean age ∼1600 BP) where as *ALDH2* does not yet appear giving insight into the strength in the southern regions and the distinct timings. A number of other possible phenotypes have been linked to these loci including reduced BMI, cardiovascular disease risk, and all-cause mortality, though the extent to which these are mediated by alcohol consumption is unclear (*35*, *36*). The concordance of the timing and geography of these events with recent evidence of the spread of tuberculosis might provide some evidence for previous theories linking these loci with mycobacterium and other pathogen resistance (*37–41*).

##### FADS1/2

We replicated signals of selection at rs174546 (C>T) in the 3’ UTR of *FADS1/2* which is associated with fatty acid metabolism, and previously reported as selected in East Asians (*17*, *21*, *42*) (*s* = 1.73% ± 0.22%; X = −6.45) (**Fig. 2.13**).This variant is not the lead variant at this locus, but is in very high LD with the lead variant (R^2^ = 0.95; D’ = 1.0). In West Eurasia, this variant begins increasing ∼9000 BP, while in East Eurasia the increase appears to start later at around ∼6000 BP (*s =* 1.52% ± 0.10%; X = 15.95).

### Alleles with evidence of parallel selection in both East and West Eurasia

#### MUC2/MUC5AC

We observed a strong signal of selection (*s* = −1.3% ± 0.16%; X = −7.00) tagged by rs12788690 (C>T) located between *MUC2* and *MUC5AC*, two mucin genes that are important components of the mucosal lining of the intestine (*MUC2*) and stomach, esophagus, and lungs (*MUC5AC*) (**Fig. 2.7**). These genes are a key component of the innate immune response but are also associated with autoimmune disorders such as Crohn’s disease and asthma, highlighting their potential for antagonistic pleiotropy. There has been one previous report of selection at an allele in moderate LD with the haplotype reported as under selection in this study in East Asia, though they did not find evidence of selection at that same variant in West Eurasia (*43*). Other mucin genes have been highlighted as selected in other populations (*44*), including this same haplotype in the ancient DNA scan in West Eurasia (*s* = −0.90% ± 0.08%.; X = −11.00).

#### IKZF3

A variant in the 3′ UTR region of the gene *IZKF3*, rs907091, is the lead SNP of a significant signal of selection in our cohort (*s* = 1.27% ± 0.16; X = 6.46) (**Fig. 2.12**). This haplotype is associated with increased risk of autoimmune disorders and asthma in the UK Biobank, Biobank Japan, and FinnGen, as well as multiple sclerosis (*45*, *46*); in Han Chinese cohorts, it is also associated with lupus (*47*). This variant is also under weaker, though still significant, selection in West Eurasia (*s* = 0.53% ± 0.08%; X = 6.46), supporting the hypothesis of contrasting disease prevalences.

#### HLA-DBQ1

As in scans of the ancient West Eurasians and in modern people (*48*), we observe strong signals of selection in the MHC region. The strongest signal is at variant rs6457619 (C>G) (*s* = −2.35% ± 0.22; X = −9.04) located in *HLA-DBQ1*, which is selected in West Eurasia as well (*s* = −0.98% ± 0.13%; X=-7.45) (**Fig. 2.1**). It is associated with increased asthma risk and elevated white blood cell counts in the UK Biobank and with rheumatoid arthritis risk in Biobank Japan. While it may still confer protection against a pathogen, today it increases risk of autoimmune disease.

### Examples of alleles with no previous evidence of selection

#### SMC5

We highlight the variant rs147421064 as it represents the only selection coefficient that is negative in East Eurasia (*s* = −2.60% ± 0.37%; X = −5.89) and positive in West Eurasia (though not genome-wide significant: *s* = 1.17% ± 0.38%; X = 3.10) (**Fig. 2.19**). It has no known phenotypic associations, but is located within *MAMDC2-AS1*, implicated in herpes simplex virus 1 infection (*49*).

#### DENDD1A

rs144867188 (T>G) in an intron of this gene has *s* = −1.55% ± 0.26% (X = 5.91) over the past 6000 years (**Fig. 2.38**). One previous study reported selection at this gene though the tagged haplotype is only in moderate LD (R^2^ = 0.10; D′ = 0.9) with our lead SNP, and is associated with an increased risk of polycystic ovarian syndrome (*50*). This variant is present at low frequencies (>4%) in West Eurasia and is not under selection there with its frequency increasing over the last 10000 years likely due to admixture or drift.

#### NRG1

An intronic variant in this gene rs2976530 (T>C) is under selection in our cohort (*s* = 1.35% ± 0.20%; X = 5.51) (**Fig. 2.39**). *NRG1* has been implicated in long-range haplotype tests using modern data (*17*), but the associated tag SNP (rs4489283) in that study is unlinked to the variant detected here (R^2^=0.00; D′=0.01).

#### CYP24A1

A haplotype upstream of this gene, tagged by rs12624784 (C>T), is under selection in East Eurasia (*s* = 1.87% ± 0.27%; X = 5.80) but the signal is largely absent in West Eurasia (**Fig. 2.25**). This variant is associated with albumin/globulin ratio in Biobank Japan, and is involved in vitamin D metabolism (*51*), a pathway implicated in selection in West Eurasia though not at this gene.

#### HIVEP3/GUCA2B

A locus upstream of both genes is tagged by rs12565534 (T>C) (*s* = 1.28% ± 0.16%; X = 6.46) and is associated with increased eosinophil count and pulse pressure in the UK Biobank (*52*) (**Fig. 2.11**). *HIVEP3* is involved in immune cell counts, and *GUCA2B* in salt homeostasis particularly in the intestines. The lead variant is in LD (R^2^ = 0.61; D′ = 0.87) with a variant implicated in hypertension in a modern Japanese cohort (*53*).

## Extraordinary correlation between East and West Eurasian selection

The most striking pattern in our data is that many of the lead variants in East Asia also have significant evidence of selection in West Eurasia.

To quantify this finding, we compared the results between these two scans, focusing on selection coefficients from the top genome-wide significant results in both East and West Eurasia. The 40 significant loci in the East Eurasian scan have a Pearson’s correlation of 0.67 (*P* = 1.98e-6) with the selection coefficients at the same variants in West Eurasia (**Fig. 3A**). For the 479 significant hits reported for West Eurasia, the measured selection coefficients in East Eurasia have a Pearson’s correlation of 0.68 (*P* = 1.98e-44) (**Fig. 3B**). A number of our independent, lead variants (13 identified in East Eurasia and 7 identified in West Eurasia) also reached significant X scores in both cohorts. This is in line with similar findings in ref. (*7*) for a set of 31 loci obtained from 5 global populations.

**Figure 3.**
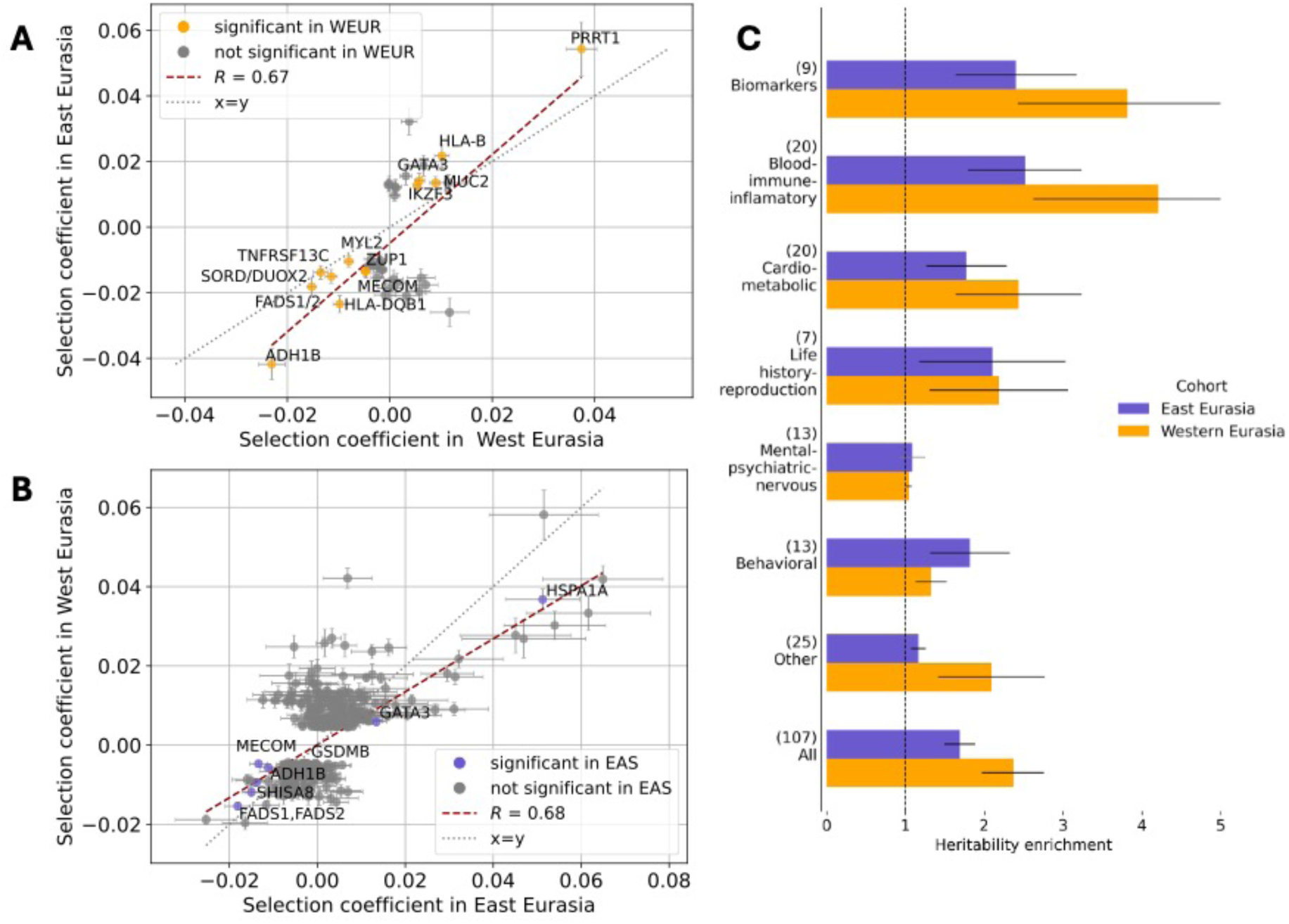
Selection scans reveal correlation of signals between West and East Eurasia. **(A)** Strong correlation between the 40 significant signals of selection in the East Eurasian cohort (EAS) and in the West Eurasian cohort (WEUR) of ref. (*1*) was observed. Those in orange where significant in both cohorts. **(B)** Strong correlation is observed between the 479 significant signals of selection in Akbari et al. and selection coefficients estimated for those same variants in the East Eurasian cohort. **(C)** S-LDSC estimates the heritability enrichment of the selection annotation of the top 1% most significant variants in our scan and in ref. (*1*) for several meta-categories of traits. We observe similar patterns of functional enrichment in both cohorts.

As a complementary way to explore the extent of these similarities, we examined combinations of variants that predict traits in genome-wide association studies and evaluated the extent to which they are similar or different in their trajectories over time in East and West Eurasia. We computed the same three tests for polygenic selection reported in ref. (*1*), γ, γ_sign_, and r_s_, which compare the polygenic score change, the signed polygenic score change without magnitudes, and the genetic correlation for each trait (**Table S4**). Using allelic effect weights measured in the UK Biobank, we observe a high correlation of the three tests both within East Eurasia (*r >* 0.63) (**Fig. 4A-C**) and within West Eurasia for the same tests (*r* > 0.67) (**Fig. 4D-F**). Repeating this analysis using allele effects weights measured in East Asian GWAS (BBJ and TPMI), we also demonstrate correlation, especially when we restrict to more stringent p-value thresholds to calculate the polygenic scores, which has the effect of reducing noise (BBJ: *r_γ_* = 0.55, *r_γ_*_-sign_ = 0.54; TPMI: *r_γ_* = 0.55, *r_γ_*_-sign_=0.48; with thresholding at genome-wide significance *p <* 5e-8) (**Supplementary Text E; Fig. E.1,E.2**).

**Figure 4.**
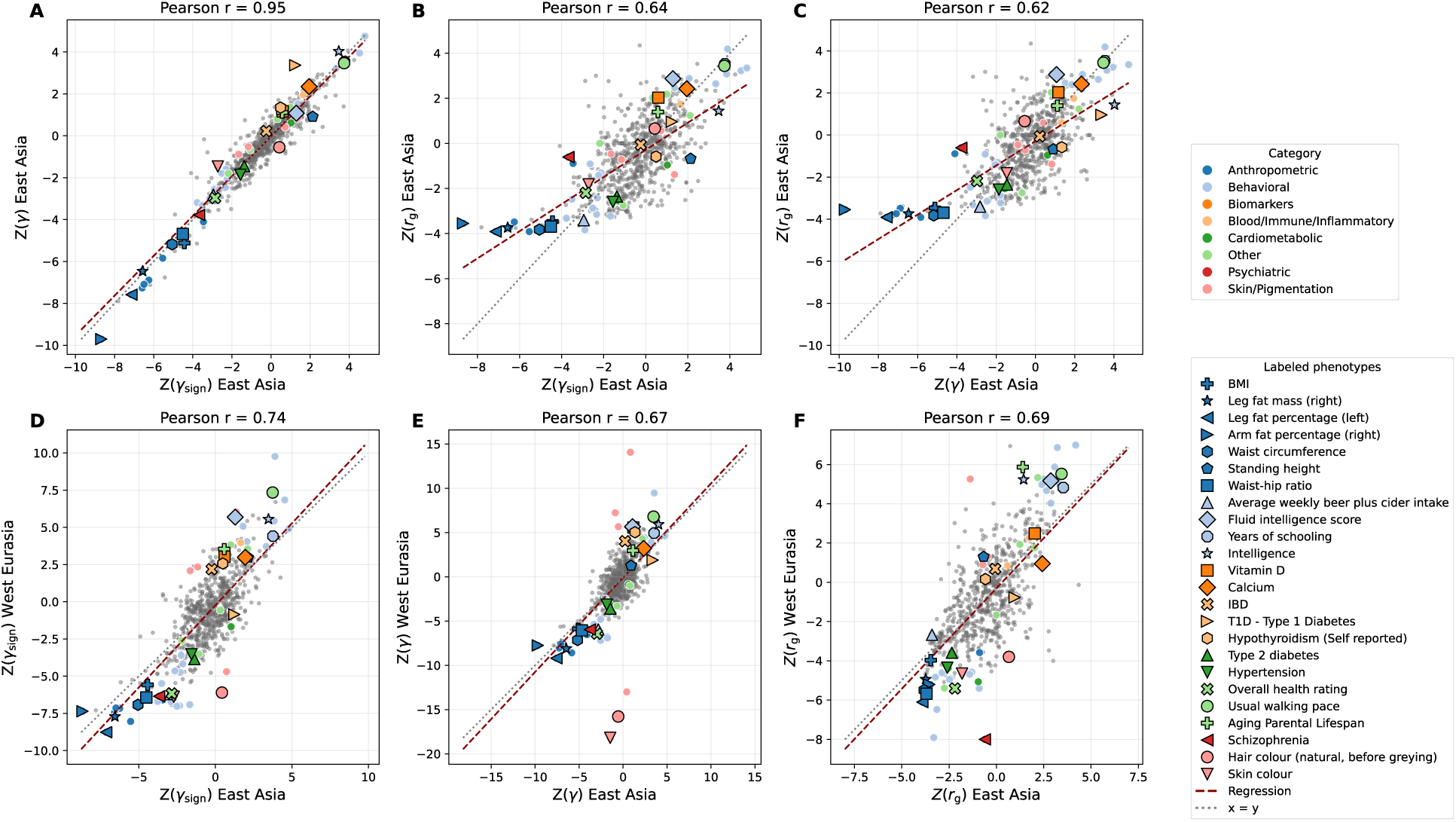
Consistency in polygenic tests of selection. (**A-C**) There is strong correlation between the 3 tests of selection from ref. (*1*) in the East Eurasian cohort including γ, γ_sign_, and rg. **(D-F)** There is correlation between each of the polygenic tests for selection in the East Eurasian and West Eurasian cohort for the same phenotypes.

As an example of correlated polygenic signals, we highlight traits associated with body fat percentage, with strong signals of selection in both cohorts, using effect weights learned from people of European ancestry in the UK Biobank. We further observed relatively strong polygenic selection in other traits whose polygenic score is correlated with BMI (*1*), often with qualitatively quite different phenotypes (e.g. years of schooling). These point to a likely shared selection pressure operating on this suite of correlated traits though the underlying biological trait driving selection remains unidentified. We also note a correlation between traits measured in both the UK Biobank and BBJ within East Eurasia (**Fig. S1**). But not all traits were correlated. We failed to observe strong selection at skin pigmentation in the East Eurasian cohort even though this is the strongest signal in West Eurasia. While the polygenic scores (PGS) used to generate the statistics shown in **Fig. 4** were calculated from GWAS of individuals primarily of European ancestry and many of the major skin pigmentation alleles are known to be different between those of European and East Asian ancestry (*54*), in ref. (*1*) the strong signal of polygenic selection persisted even after dropping the top strongest-associated loci from the analysis. To investigate this further, we tested polygenic scores from ref. (*55*) based on East Asian cohorts; however, we observe no significant result after correcting for multiple hypothesis testing for γ or γ_sign_ at any p-value thresholding (-log_10_ (*P*) < 2.0-7.3) (**Table S5**). Taken together with the lack of single-variant results associated with pigmentation, our analyses suggest that the timing of pigmentation adaptation is substantially different from that observed in West Eurasia in line with findings reported in a recent global selection scan (*7*). While strong depigmentation occurred in West Eurasia in the last ten millennia, our results show that genetic selection for skin lightening in East Asia preceded the Holocene.

We finally observe an extraordinary similarity in the functional categories of variants that are enriched for selection in East and West Eurasia. We used Stratified LD Score Regression (S-LDSC) to measure the heritability enrichment of our selection annotation of the top 1% of variants based on significance for different categories of GWAS traits (**Materials and Methods**). We detected enrichment in blood/immune/inflammatory, cardiometabolic, life history/reproduction, mental/psychiatric/nervous, and behavioral traits; the ordering of trait effects was qualitatively similar to the ordering of effect sizes as in West Eurasia (**Fig. 3C**).

## Discussion

The sample size we analyze is an order of magnitude less than the West Eurasian scan (*1*), and the samples are far more diverse in terms of their geography and economies (for example, we include steppe pastoralists, farmers in the East Asian core region, and Oceanians living in Pacific islands). This limits statistical power, likely explaining the reduced number of genome-wide significant loci (40 versus 479). Increasing sample size by an order of magnitude would plausibly yield a similar number of genome-wide discoveries as in West Eurasia, providing a rich basis for understanding the extent to which the effect of selection on the genetic architecture of complex traits in East Asia is the same or different from that in West Eurasia.

Our finding of highly correlated selection signals in East and West Eurasia is extraordinary in light of the fact that the two cohorts have had separate histories since they diverged. While about a third of the ancestry of our cohort is derived from West Eurasians mostly Bronze Age steppe pastoralists who migrated eastward in the second and first millennia BCE, the correlated signals cannot be reflecting this shared ancestry: simulations show that the GLMM method only detects selection occurring in the time series itself, not in the populations whose descendants migrated to the region (*1*). To further check the robustness of this finding, we repeated the analysis restricting to individuals with relatively high East Asian ancestry (>80% as represented by the 1000 Genomes JPT population in supervised ADMIXTURE), and we observe a high correlation between the single-variant and polygenic tests for selection with the West Eurasian scan (**Fig. S2,S3; Supplementary Text C).**

Our study shows that similar lifestyle changes at the two ends of Eurasia—catalyzed by the advent of agriculture, the spread of pastoralism, and exposure to new types of pathogens as populations grew and people began living closer to each other and to their animals—produced similar genetic effects in populations with independent histories. Both West and East Eurasians derive their genetic variation from a shared ancestral population that spread out of Africa and the Near East after around 50,000 years ago. The frequencies of variants in that ancestral population were shaped by patterns of selection in hunter-gatherer groups. But our results suggest that once exposed to agriculture, pastoralism, modern pathogens, and the lifestyles that come with living in larger state societies, the variants inherited from this ancestral population continued moving in the same direction in both West and East Eurasia, likely reflecting convergent evolution to adapt to the new requirements of food producing lifestyles and living at high-density. An important topic for future research will be to carry well-powered time-transect studies in other world regions and times with similar environment exposures, for example Native Americans in the Central Andes or in Mesoamerica who have had similar exposure over thousands of years to the transition to food production. Equally illuminating would be time-transect studies in foragers, for example, Native Americans in the southern cone of South America, or hunter-gatherers in both Europe and East Asia prior to the transition to farmers, once sufficient sample sizes become available. Our results raise the possibility that selection on many of these traits is due to the transition to food production; time series in forager populations would make it possible to test this hypothesis. Comparing and contrasting trajectories global will result in better understanding of the selective pressures such as pathogen exposures and lifestyle transitions shaping our genome and allow insights into human health today.

## Acknowledgments

We are grateful to Tianyi Wang, Daniel Tabin, and Yue-Chen Liu for helpful conversations and suggestions for this project. We thank Nicole Adamski, Rebecca Bernardos, Nasreen Broomandkhoshbacht, Kim Callan, Alex Claxton, Olivia Cheronet, Elizabeth Curtis, Matthew Ferry, Trudi Frost, Ilana Greenslade, Eadaoin Harney, Lora Iliev, Aisling Kearns, Jack Kellogg, Ann Marie Lawson, Megan Michel, Jonas Oppenheimer, Iris Patterson, Susanne Nordenfelt, Lijun Qiu, Kristin Stewardson, Anna Szécsényi-Nagy, H. Noah Workman and Fatma Zalzala for support in the wet laboratory; and Iosif Lazaridis, Heng Li, Adam Micco, Mariam Nawaz, Zhao Zhang, and Mengyao Zhao for contributions to the bioinformatic processing. We are also grateful to the more than 80 archaeologists and anthropologists who provided explicit permission for release of raw data for individuals with never-before-reported ancient DNA data they shared with us for the purposes of studies of selection, decoupled from the contextual information which will be presented in papers on which they are co-authors and is necessary for any studies of the population history of these individuals. Finally, we would like to acknowledge the individuals both living and deceased whose contributions made this work possible.

## Funding

Getty Foundation Dual Postdoctoral Program (ARB).

National Institutes of Health grant HG012287 (DR).

Allen Discovery Center program, a Paul G. Allen Frontiers Group advised program of the Allen Family Philanthropies (DR).

John Templeton Foundation grant 61220 (DR).

Private gift by Jean-Francois Clin (DR).

Howard Hughes Medical Institute (HHMI). (DR).

## Author contributions

Conceptualization: ARB, AA, DR

Methodology: ARB, AA

Investigation: ARB, AA, DR

Funding acquisition: DR

Sample and Sequencing processing: NR

Bioinformatic processing: SM

Sample coordination and preparation: RP

Project administration: DR, RP, NR, SM

Supervision: AA, DR

Writing – original draft: ARB, DR

Writing – review & editing: ARB, AA, DR, NR, SM, RP

## Data, code, and materials availability

The software used to run the GLMM analysis, PQLseqPy, is available from https://github.com/mokar2001/PQLseqPy. All other tools are noted in the methods where appropriate

**Data S1** and **Data S2**, which include the summary statistics for the GLMM analysis, will be available upon final publication. For earlier access, please write to the corresponding authors.

Aligned sequences for the newly reported data from 867 ancient individuals are available through the European Nucleotide Archive under an accession number that will be made available upon final publication. This includes complete genetic data for all newly reported individuals alongside point estimates of their dates and geographic/genetic clustering into three regions of East Eurasia, which is the full data used in the present study. Our release of these data without restrictions to enable studies of natural selection has written approval from third-party sample custodians. Please contact corresponding author DR for any questions regarding metadata not used in this study—such as skeletal codes, latitudes and longitudes, site names, site descriptions, physical anthropology, and cultural context—which will be reported in future work that should be the references for studies of population history and archaeology. Imputed genomes for ancient individuals are available at the Dataverse repository at an accession number that will made available upon final publication.

## Supplementary Materials

Materials and Methods

Supplementary Text

Figs. S1 to S3

Tables S1 to S6

References (*1–70*)

