## Supplementary Materials for "Convergent natural selection at both ends of Eurasia during parallel radical lifestyle shifts in the last ten millennia"

**The PDF file includes:**

Materials and Methods
Supplementary Text
Figs. S1 to S3
References

### Materials and Methods

aDNA Data Generation

Wet laboratory and bioinformatic processing of newly reported ancient DNA sequences follow the methods outlined in ref. (*1*).

Sample Quality Control

We filtered to ancient samples by generating a PCA based on modern samples and selecting those that clustered together in PC space (ensuring minimal West Eurasian ancestry) (**Supplementary Text A**). We further restricted to individuals sampled east of 65°E longitude to avoid geographic overlap with ref. (*1*) which was restricted to samples five degrees to the west of this line. Modern samples were represented with whole genome sequencing data: 504 East Asians from the 1000 Genomes Project Phase 3 (*56*) (all in the CHB, CHS, CDX, KVT, and JPT cohorts) and 106 from the Human Genome Diversity Project and other publications. HGDP samples and other moderns not from 1000 Genomes Project were imputed with the same pipeline as ancient samples. 1000 Genomes Project samples were not imputed as they made up the reference panel. We implemented all additional sample quality control as in ref. (*1*) with one exception (**Supplementary Text A**): we did not remove relatives from the selection scan as the genetic-relatedness matrix can correct for this bias. (This was not performed in ref. (*1*) due to the sufficient sample size and to increase the speed of the analysis).

Imputation

Imputation followed ref. (*1*) using GLIMPSE (v1.0.0) (*57*) after using bcftools mpileup (v1.13) (*58*) to generate genotype likelihoods for all SNPs and indels. The high coverage (30x) 1000 Genomes Project phase 3 (*56*) sequences were used as a reference panel after using CrossMap (v0.5.2) (*59*) to convert to GRCh37/hg19. Genotype likelihoods for indels were set to missing prior to imputation.

Variant Quality Control

We started with 9.7 million variants that passed quality control metrics for ref. (*1*), as these had been assessed for batch effects between various technologies including between capture, shotgun, and modern sequencing data. We further filtered within our East Eurasian dataset to remove variants with poor imputation or batch effects that were too rare to be filtered in the West Eurasian quality controls (**Supplementary Text B**). We added additional filters for INFO score greater than 0.6 on average for both the captured and shotgun samples in our East Asian dataset. Additionally, using 114 samples from this study for which we had both shotgun and capture data, we calculated an error rate for each variant comparing the sequences from the same individuals and filtered those under a threshold as in ref. (*1*). Following ref. (*1*) we generated ancestry matched sets of ancient and modern samples to assess batch effects (**Supplementary Text B**). This left 7.5 million variants. Finally, using two runs of the GLMM with and without potential batch effect related covariates (imputation quality score, sequencing technology, ancient status, and 1kg membership) we filtered variants with a residual of the regression of these runs greater than 3 (**Supplementary Text B**). This left 7,149,968 well-imputed variants for downstream analysis including 6,379,693 SNPs and 770,275 indels with non-zero frequencies. This resulted in summary statistics at 6,079,037 variants after requiring a MAF of 0.5%, a MAC of 50, and convergence of the GLMM model in all runs. All variants are reported with the selection coefficient as on the derived allele in the text and as at the minor allele in GNOMAD (*60*) (**Data S1**).

Generalized Linear Mixed Model

We employed the test for directional selection presented in ref. (*1*). To increase our sample size, we retained relatives, which changed our clustering strategy such that all individuals with an up to 2$^{nd}$ degree relative in the dataset were assigned their own cluster and only unrelated samples were assigned to clusters resulting in 500 multi-individual clusters and 870 total clusters. This included running a model with only the genetic relatedness matrix as a covariate and running a model with additional covariates for sample imputation quality, sequencing technology (capture versus whole-genome sequencing) for ancient data, ancient status of sample, and modern status of sample. The z-scores between both runs were regressed against each other as in ref. (*1*) with some modifications based on the properties of this dataset (**Supplementary Text B**), and those with a residual larger than 3 were removed from the scan. Due to the power, we also required a minimum MAF in the whole cohort of 0.5% and a MAC of 50. To determine the threshold for significance, we adopted an approach validated in simulations in ref. (*1*) whereby we find a plateau of enrichment using functional annotations. In addition to the Pan-UKB (*52*) and GTEx (*9*, *10*) eQTLs and sQTLs utilized in ref. (*1*), we added annotation from the Biobank of Japan (BBJ) (*11*) and the Taiwan Precision Medicine Initiative (*12*) (TPMI) (**Fig. 1B**), hypothesizing that they might be more relevant to our cohort of predominantly East Eurasian ancestry. From this enrichment, we determined a significance threshold of $P < 5.75E\text{-}11$, at which the enrichment of the combined BBJ and TPMI genome-wide significant hits is 8-fold that of all tested variants and that using the UKBB is 3.2-fold, which we adjust to the selection statistic in ref. (*1*) taking Z/1.20 (the equivalent Z score is 6.55, 1.20-fold that of the genome-wide significance threshold at 5.45). The smaller sample size of our cohort compared to ref. (*1*) (~8x smaller) resulted in less power to filter variants based on batch effect particularly when utilizing the matched modern and recent (>1000 BP) samples and ancients for which both shotgun and capture data was available. This was particularly true of variants common in East Eurasia, but rare or absent in West Eurasia. Thus, to be more conservative, and follow GWAS approaches, we required at least two variants in a region to pass the significance threshold in order to consider a single locus under directional selection. We fine-mapped variants and arrived at our total number of independent loci under selection by controlling for the linkage disequilibrium within our cohort, taking a conservative approach to independence of $R^2 < 0.05$ for each block and DP < 0.8.

Polygenic Analysis

We implemented tests for polygenic selection as presented in ref. (*1*). This included both the $\gamma$ and $\gamma_{sign}$ tests based on change in mean polygenic score over time using GEMMA (v0.98.5) (*61*) and a GRM to again account for population structure as well as $r_s$, the genetic correlation as calculated using LDSC (*62*, *62*, *63*) or S-LDXR (*64*) depending on whether the original GWAS was performed in the same (East Asian) or different ancestry (West Eurasian) populations respectively (**Table S4**). We tested all phenotypes included in ref. (*1*).

East Asian ancestry analysis

To determine the extent to which our results might be biased by the influence of West Eurasian ancestry in the region, we restricted to 1361 individuals with a high proportion of East Asian ancestry as represented by the 1000 Genomes JPT population in supervised ADMIXTURE (*65*, *66*), competing with the GBR population representing West Eurasian ancestry. We reran the single variant analysis following the same procedure as the main cohort (**Data S2**). We compared the correlation of the selection coefficients as in **Fig. 3A,B** of the 40 variants significant in the original East Eurasian analysis (one variant dropped out due to low minor allele count in the restricted cohort) and the 239 significant variants reported in ref. (*1*) with MAC > 10 in the restricted cohort. Both retained a high correlation $r = 0.77$ ($P < 1.30\text{e-}8$) for the East Eurasian hits and $r = 0.60$ ($P < 1.75\text{e-}29$) for the West Eurasian hits (**Fig. S2**). We similarly repeated the polygenic analysis with this smaller cohort and found $r > 0.59$ for the three tests (**Table S6; Fig. S3**).

East Asian Summary Statistics

Summary statistics for GWAS from Biobank Japan were downloaded from: https://pheweb.jp/downloads. Summary statistics from the Taiwan Precision Medicine Initiative were obtained from: https://pheweb.ibms.sinica.edu.tw/phenotypes. We included all phenotypes for which there was at least one genome-wide significant hit. TPMI associations were lifted over to GRCh37 using rsIDs. $R^2$ values reported for modern East Asian and European data were calculated using LDlink (ldlink.nih.gov)

### Supplementary Text

#### A. Sample Quality Control

Sample quality control followed the approach outlined in ref. (*1*). We selected samples based on PCA coordinates and geographic restrictions. Modern samples were drawn from the 1000 Genomes Project Phase 3 and included all samples in the CHB, CDX, KVT, and JPT cohorts (N=504) and 106 individuals from the Human Genome Diversity Project and other publications with whole-genome sequencing. Modern samples not from 1000 Genomes Project were imputed with the same pipeline as ancient samples. 1000 Genomes Project (*56*) samples were not imputed as they made up the reference panel.

We implemented additional sample quality control as described in ref. (*1*) was implemented however relatives were not removed as the smaller sample size allowed for them to be included as independent clusters in the GLMM (**Table SA.1**). This included restriction to samples with a mean imputation quality score (IQS) > 0.9 and little evidence of contamination as defined in ref. (*1*), leaving 2,472 individuals (**Table SA.2**).

**Table SA.1:** Quality control filters and remaining samples at each step.

| Filter | Remaining samples | Notes |
|---|---|---|
| Filter 1 | 2,820 | Restrict to Eastern Eurasian ancestry using PCA. |
| Filter 2 | 2,511 | Remove sequences with evidence of contamination or damage, abnormal sex ratios, unreliable estimates, or poor dating. |
| Filter 3 | 2,511 | Remove sequences with too many relatives (> 100). |
| Filter 4 | 2,510 | Remove sequences if the dates of their first- or second-degree relatives vary widely, defined as a standard deviation > 400 years. |
| Filter 5 | 2,220 | Remove identical sequences. |

| | | |
|---|---|---|
| Filter 6 | 2,219 | Remove sequences with abnormal chromosome coverages. |
| Fitler 7 | 2,202 | Remove sequences whose date is not predicted by PC regression (**Fig. SA.1**) |
| Filter 8 | 2,114 | Remove individuals with close relatives who have discordant predicted dates. |
| Filter 9 | 1,968 | Remove overlap with any iteration of ref. (*1*) analysis. |

Filters that varied from ref. (*1*) are explained as follows: **Filter 2: Restrict to East Eurasian ancestry.** After performing agglomerative clustering on the top 100 PCs, we applied the following restrictions to mean PC values for each cluster: PC1 > 0.081 and -0.04 < PC3 < 0.04. Sequences in clusters that met these criteria were included (**Fig. SA.1**). We further removed any individual samples west of 65°E longitude. This left us with 1,968 individuals which were combined with the 504 1000 Genomes East Asian samples (CHB, CHS, CDX, KVT, JPT).

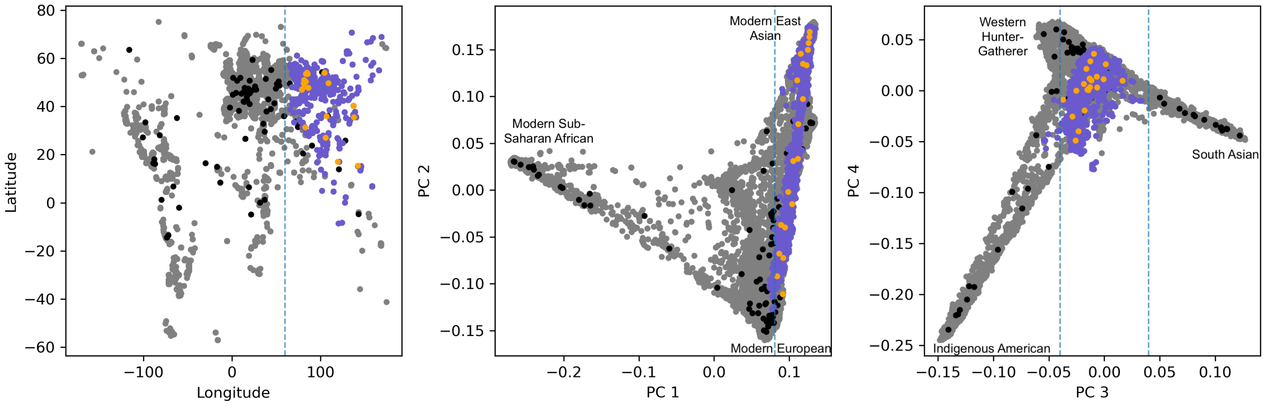


**Fig. SA.1:** Purple circles represent the individuals included for the first step in our sample QC and gray circles represent excluded individuals. Orange circles show the mean values of included clusters and black circles show the mean values of the excluded clusters. Blue dashed lines on each axis show the criteria applied for inclusion.

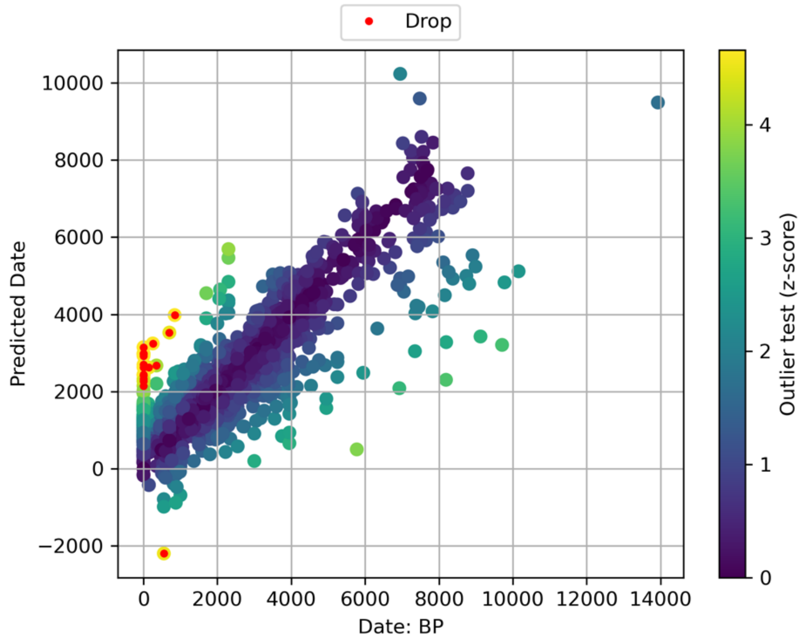


**Figure SA.2:** Predicted date based on the top 100 PCs versus the reported date for samples. Outliers highlighted in red were dropped from the analysis.

**Table SA.2:** Individuals analyzed in this study.

| Individuals | Description |
|---|---|
| 1,605 | **1. Previously published individual (no newly reported data)**<br>610 present-day (610 shotgun)<br>995 ancient (651 capture, 344 shotgun) |
| 12 | **2. Previously published individual with new shotgun sequences** |
| 38 | **3. Previously published individual with new capture data** |
| 817 | **4. Never-before reported individual**<br>813 capture, 4 shotgun |

B. Variant Quality Control

The pipeline for variant quality control utilized the approach in ref. (*1*) , including filtering all low-quality variants as called in that dataset. We generated several subsets of data to check for allele frequency consistency between datasets as outlined in **Table SB.1**. As in Step 2 of variant QC in ref. (*1*), we created genetically matched cohorts between 1KG and our recent ancient samples to test for batch effects by using hierarchical clustering of the top 10 PCs into 25 clusters (**Fig. SB.1**).

**Table SB.1:** Sample set used for variant quality control.

| Sample set name | Number of samples | Description |
|---|---|---|
| aDNA_SG | 103 | Shotgun sequences of ancient individuals with both shotgun and 1240k enrichment reagent data |
| aDNA_1240k | 103 | 1240k sequences of ancient individuals with both shotgun and 1240k enrichment reagent data |
| EAS1 | 421 | Eastern Eurasian individuals in our dataset with 0<Date<1k years BP |
| EAS0 | 105 | Eastern Eurasian individuals in our dataset with Date = 0 years BP but not in 1KG |
| GNOMAD_EAS | 780 | Allele frequencies in the East Asian subset of GNOMAD are presented in the gnomad_genomes_an_EAS column of the variant manifest of pan-ukbb (2022-04-11). |
| 1KG_ALL | 2504 | All the individuals from 1000 GP |
| 1KG_EAS | 504 | 1000 GP individuals from EAS populations |
| 1KG_CN | 301 | 1000 GP individuals from CHB, CDX, and CHS populations |
| EAS0_match_1KG | 31 | 1KG individuals that are ancestry matched with individuals from EAS0 |
| 1KG_match_EAS0 | 31 | EAS0 individuals that are ancestry matched with individuals from 1KG |
| EAS1_match_1KG | 44 | 1KG individuals that are ancestry matched with individuals from EAS1 |
| 1KG_match_EAS1 | 44 | EAS1 individuals that are ancestry matched with individuals from 1KG |
| EAS0_match_1KG_CN | 21 | EAS0 individuals that are ancestry matched with individuals from 1KG_CN |
| 1KG_CN_match_ EAS0 | 21 | 1KG_CN individuals that are ancestry matched with individuals from EAS0 |
| EAS1_match_1KG_CN | 32 | EAS1 individuals that are ancestry matched with individuals from 1KG_CN |
| 1KG_CN_match_ EAS1 | 32 | 1KG_CN individuals that are ancestry matched with individuals from EAS1 |

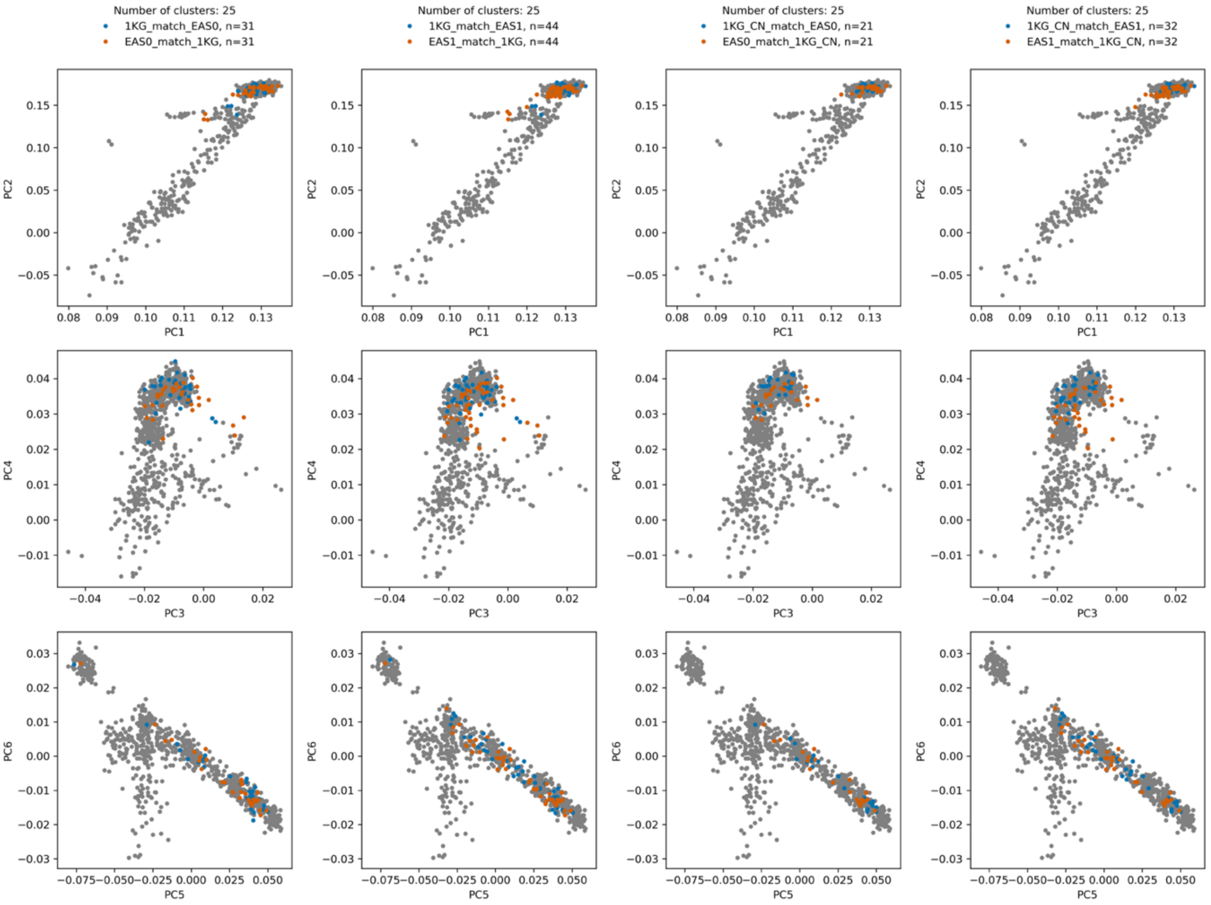


**Figure SB.1:** The top 6 PCs for samples matched by across different datasets. In the hierarchical clustering procedure, the n_clusters parameter is set to the number of clusters labelled for each set; the final number of ancestry-matched individuals in each set is indicated by n.

As in ref. (*1*), for individual *i* with both shotgun and 1240k sequences that are in the aDNA_SG and aDNA_1240k sets of sequences, we define $\Delta_i$= ($GT_i$ (Shotgun) – $GT_i$ (1240k))/2 for each variant such that $GT_i$ (Y) means the imputed genotype in data type Y (shotgun or 1240k) for individual *i* for the variant of interest. A variant must meet the following criteria to pass QC:

1. Minor allele frequency in 1KG_ALL > 0.005
2. mean (|Δ|)<0.05
3. The p-value of the null hypothesis mean (Δ) = 0 is > 1e-5
4. P-chi2-normalized (EAS0_match_1KG, 1KG_match_ EAS0) > 1e-5
5. P-chi2-normalized (EAS1_match_1KG, 1KG_match_ EAS1) > 1e-5
6. P-chi2-normalized (EAS0_match_1KG_CN, 1KG_CN_match_ EAS0) > 1e-5
7. P-chi2-normalized (EAS1_match_1KG_CN, 1KG_CN_match_ EAS1) > 1e-5
8. INFO_EAS_1240k > 0.6
9. INFO_EAS_SG > 0.6

P-chi2-normalized (pop1, pop2) represents the p-value of the chi-square statistic divided by its mean over all variants, from the chi-square test between pop1 and pop2 given the allele counts. The consistency of the allele frequencies between pairs of ancestry-matched subsets is shown in **Figure SB.2**.

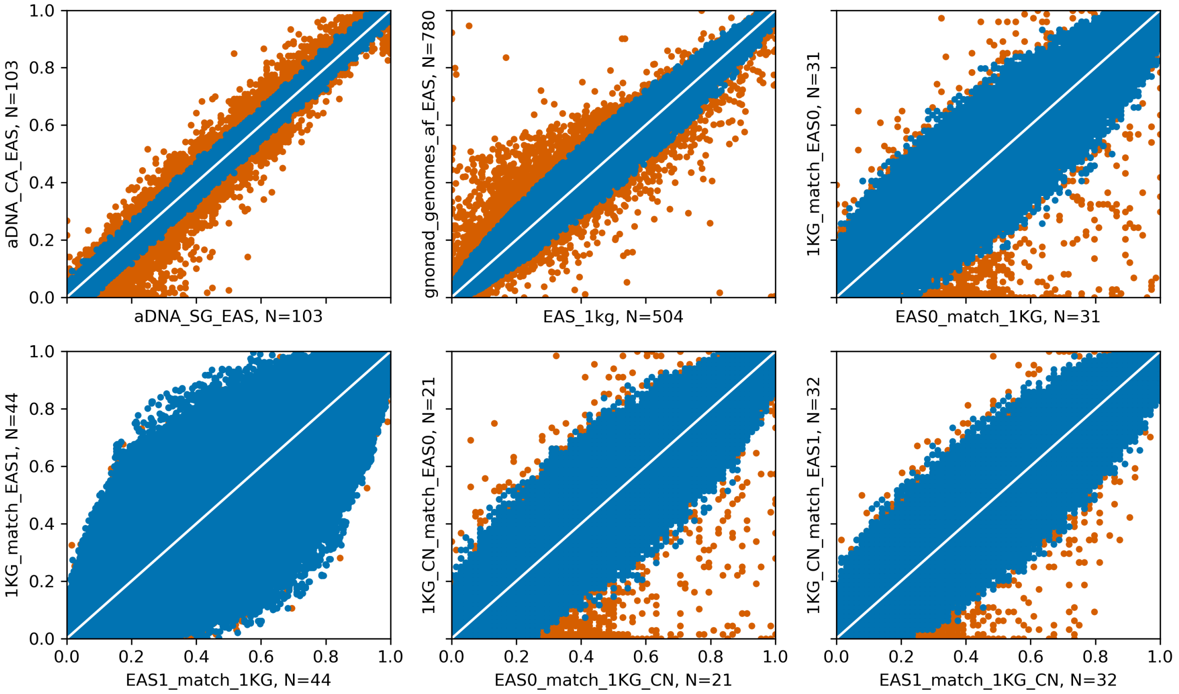


**Figure SB.2**: Allele frequency comparisons across ancestry-matched datasets. We subsample to 1,000,000 randomly selected variants that pass the filters (blue) and 1,000,000 that do not (orange) for display purposes. The total number of individuals in each set is indicated by N. The x and y axes show allele frequencies in the subset of individuals as labeled for that axis.

After running our GLMM pipeline on the remaining variants, we implemented an additional quality filter to control for several types of batch effect as in ref. (*1*) variant QC step 3. This involved running the test for selection in two rounds: one with no covariates other than the GRM and one with 4 covariates: IQS (imputation quality score), aDNA (1 for ancient individuals, 0 for modern), SG (1 for shotgun-sequenced ancient individuals, 0 otherwise), and 1kg (1 for 1000 GP samples, 0 otherwise), included as fixed effects in our GLMM. As in ref. (*1*), let $Z$ denote the Z-score of the estimated selection coefficient calculated without these covariates, and $Z_0$ the Z-score calculated with them. We calculated weighted least squares (WLS) regression for the two Z-scores and excluded all variants with Pearson residuals greater than 3 (**Fig. SB.3**).

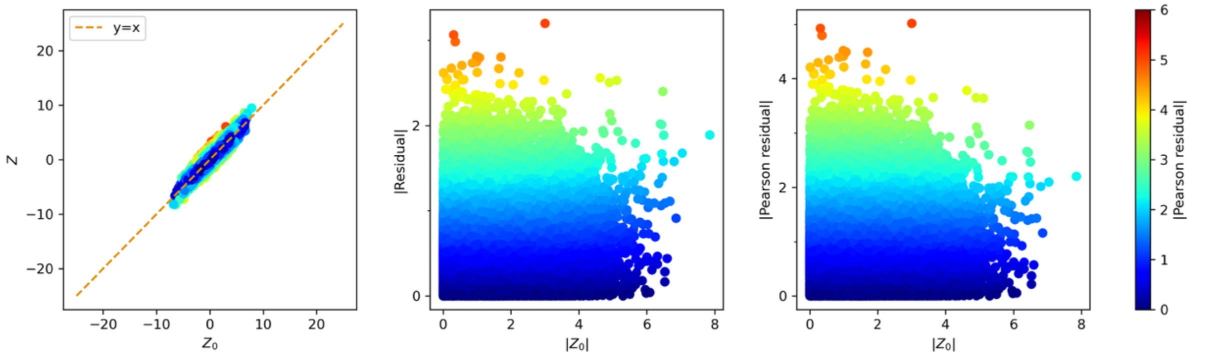


**Fig. SB.3:** Comparison of Z-scores from GLMM, with ($Z$) and without ($Z_0$) the aforementioned covariates. Outliers, which could be due to batch effects, are identified and removed using Pearson residuals from WLS. The color scale represents the |Pearson residual|.

Again following ref.(*1*), we fit a WLS for $Z = a + bZ_0$ with weights = $1/(cZ_0^3)$, such that c = 5.62-4. (**Fig. SB.4**).

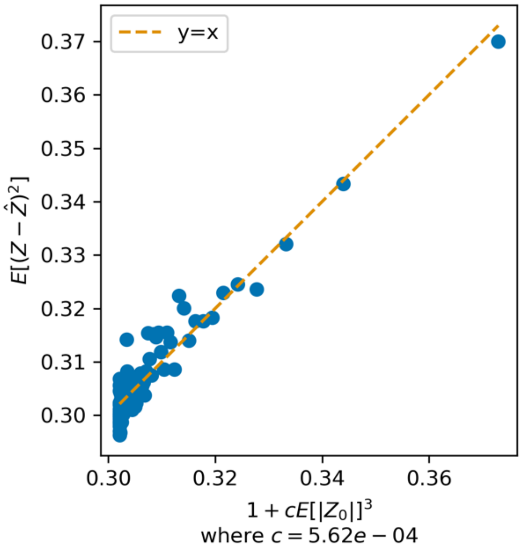


**Fig. SB.4:** Empirical inference of weights for weighted least square (WLS).

After all variant filters, 7,149,968 variants passed quality control including 6,379,693 SNPs and 770,275 indels.

| Variant set | Category based on presence in different variant sets (1: present, 0: absent) | | | Variant set count |
|---|---|---|---|---|
| SNP | 1 | 0 | 1 | 6,379,693 |
| 1240k | 0 | 0 | 1 | 853,428 |
| Category count | 5,526,265 | 770,275 | 853,428 | 7,149,968 |

**Table SB.2:** Summary of variants (SNP and indels) that passed final quality control including residual test, categorized by their presence in different variant sets

C. Ancestry composition of samples.

To assess the extent to which our samples had predominately East Eurasian or West Eurasian ancestry, we ran supervised ADMIXTURE (v1.3) (*65*, *66*) on all samples with K=2, designating the 1000 Genomes Project GBR samples as representatives for the West Eurasian source and the JPT samples as representing East Eurasian ancestry. We restricted to 1240k variants that passed our thresholds and pruned using PLINK –indep-pairwise 50 20 0.1. This provided an approximate proportion of each ancestry in our dataset as a whole and per cluster (**Fig. SC.1**).

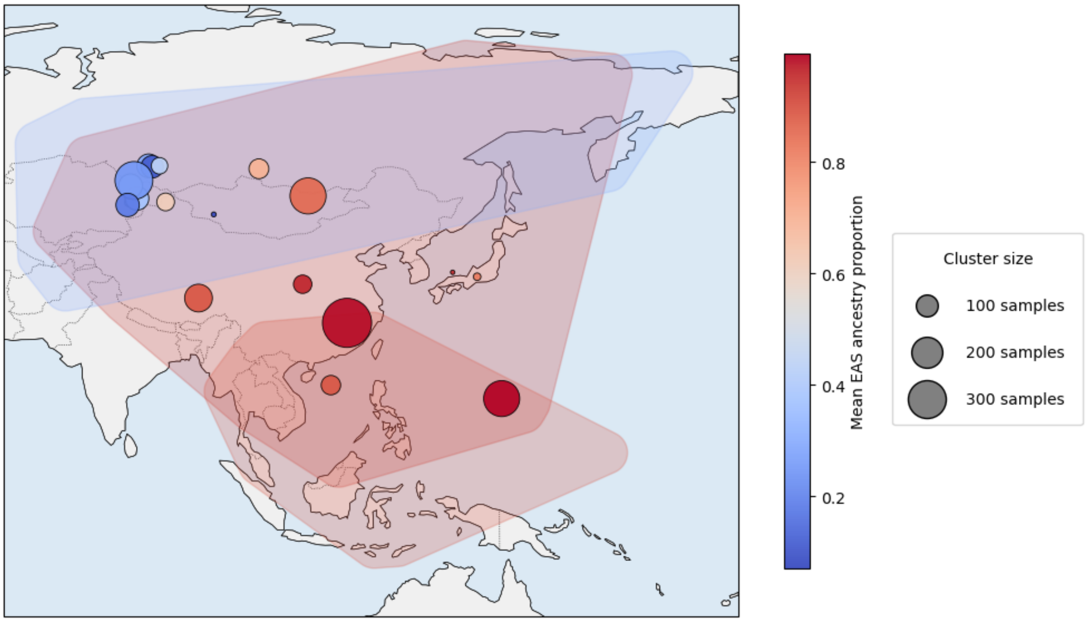


**Fig. SC.1.** Mean PCA clusters are colored by proportion of East Asian ancestry as assigned by ADMIXTURE and are sized based on number of samples. Regions overlap all samples in the assigned regions (see **Figure 1A**) and are shaded by average ancestry proportion of samples in those regions.

The Ancient South East Asian ancestry in our allele frequency trajectories (**Fig. 2**) were represented by a number of samples not used in the main analysis due to lower imputation quality, but which we included here to give an approximate estimate of the frequency of each allele in key ancestral populations. The samples were drawn from refs. (*67–69*).

D. Determining overlap with previous selection scans.

We conducted a literature review of previous studies focused on detecting selection in East Asian populations. We found 11 works with a substantial focus on global or East Asian selection scans. We did not include studies focused on selection only at specific loci, though these are cited in the text were relevant. Only 2 previous studies included ancient DNA and the remainder are focused on Fst or haplotype based methods for detection. The HLA was excluded for comparison as it was widely reported without specific haplotype information. We examined the following papers by including their results as follows:

1. Voight et al. (2006) (*14*). We took the rsIDs from Protocol S1 (Summary of Strongest Regions of Selection Genome-Wide in East Asians). One overlap was found at the *ADH1B* locus, rs980972.
2. Sabeti et al. (2007) (*15*). We used table 1 (The twenty-two strongest candidates for natural selection) and used each position with a window the size of the region on either side We lifted over coordinates from hg17 to GRCh37 using the UCSC LiftOver (*70*) web browser tool. No overlaps were detected.
3. Kimura et al. (2007) (*16*). Using data S1for the EASvsYRI and EASvsCEU tests, we converted the positions from GRCh34 to GRCh37 using the UCSC LiftOver (*70*) web

browser tool, with 18 records failing to convert. One region at *TNFRSF13C* overlapped our results.

E. Polygenic Tests for Selection

We implemented all tests for selection as in ref. (*1*). We additionally ran the $\gamma$ and $\gamma_{sign}$ tests on 220 summary statistics from Biobank Japan (*11*) (**Fig. SE.1**) and 294 summary statistics from the Taiwan Precision Medicine Initiative (*12*) (**Fig. SE.2**) to determine whether we observed the same correlation between the Z-statistics for the $\gamma$ and $\gamma_{sign}$ tests between West and East Eurasia when using summary stats ascertained on an East Asian cohort. We conducted this test using effect sizes directly as well as those th at were converted to just the sign of the effect size. We ran tests both including and excluding the MHC region and for different significance thresholds including -log(P) >2.0, 3.0, 4.0, 5.0, 6.0, 7.0, and 7.3. Comparing across these categories demonstrated that the correlation in these summary statistics increased with increasing significance thresholds for included variants. In contrast to the summary statistics for West Eurasia, which are primarily derived from the UK Biobank, the correlation is relatively weaker for the same significance thresholds. However, this is likely due to the overall reduced power of both biobanks based on sample size and as previously demonstrated by wider variance in the heritability in Biobank Japan (*64*).

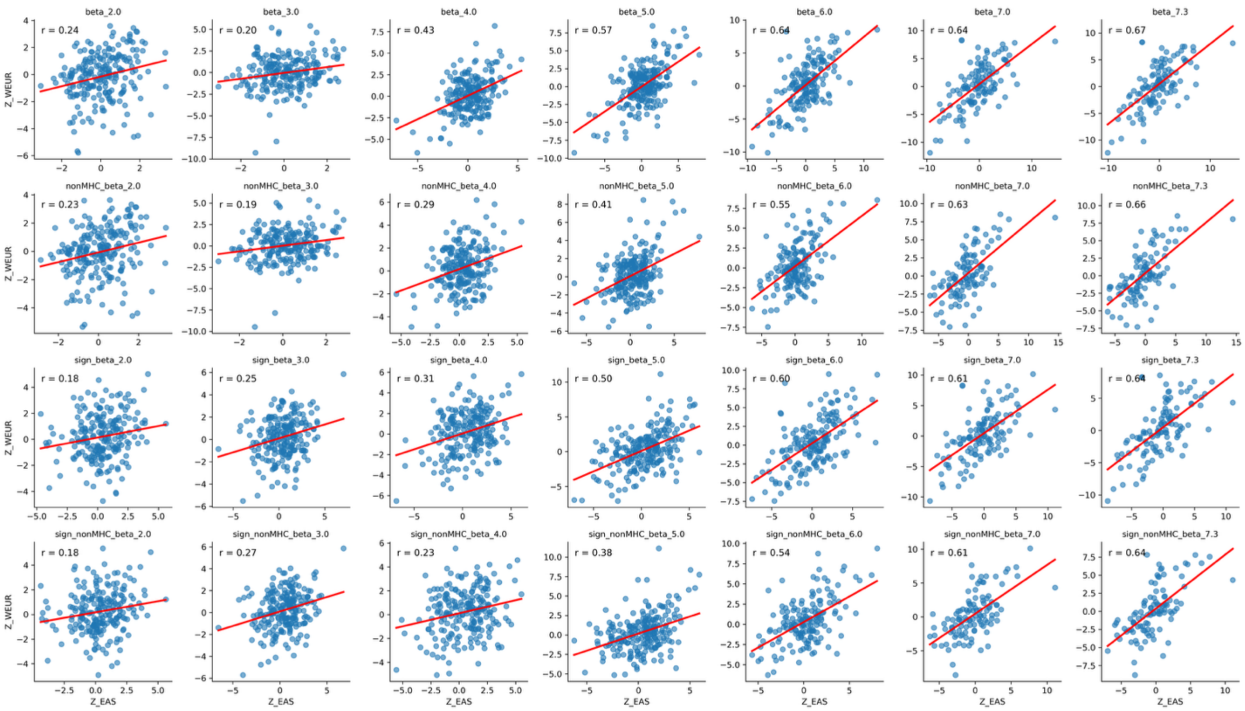


**Figure SE.1.** Polygenic tests for directional selection in West Eurasian and East Eurasian cohorts using summary stats from Biobank Japan.

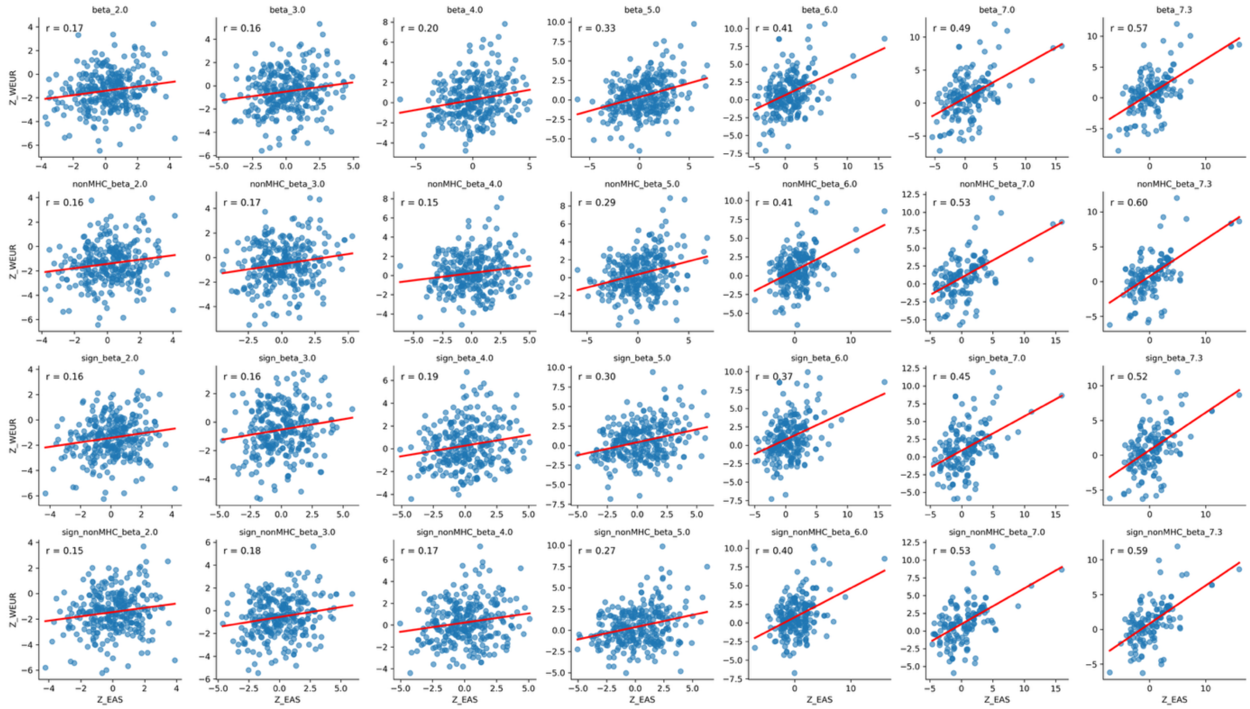


**Figure SE.2.** Polygenic tests for directional selection in West Eurasian and East Eurasian cohorts using summary stats from Taiwan Precision Medicine Initiative.

Skin pigmentation has been shown to have different genetic architectures in different populations. To confirm that the lack of polygenic selection we observed on pigmentation traits based on GWAS calculated from the UK Biobank was not in fact due to this ancestry difference, we also tested three traits from ref. (*55*) via the GWAS catalog, which were ascertained in East Asians for the following phenotypes: skin luminance (GCST90320257), and red-green skin component (GCST90320257), and yellow-blue skin component (GCST90320259) (**Table S5**).

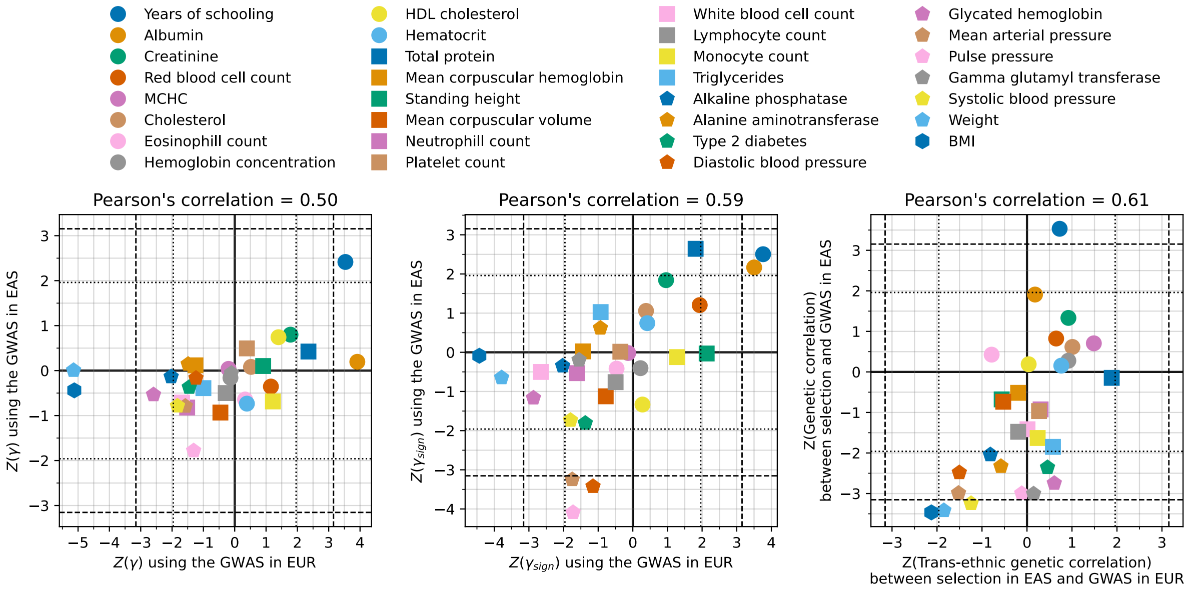


**Fig. S1.**

For a subset of phenotypes that have independent summary stats ascertained in both an East Asian (EAS; primarily BBJ) and West Eurasian (EUR; UKB) GWAS, we observe correlation in the test statistics for the East Eurasian cohort.

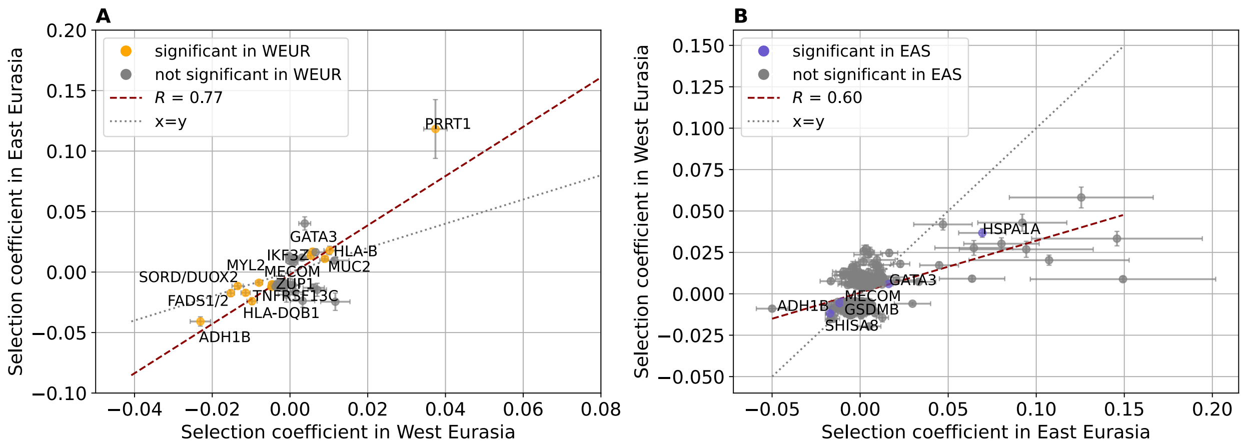


**Fig. S2.**

Correlation in selection coefficients is maintained in a high East Asian ancestry cohort. **(A)** Strong correlation between the 40 significant signals of selection in the East Eurasian cohort and in the West Eurasian cohort of ref. (*1*) was observed. One variant dropped out due to low minor allele count in this subset of the cohort. Those in orange where significant in both cohorts. **(B)** Strong correlation is observed between the 479 significant signals of selection in ref. (*1*) and selection coefficients estimated for those same variants in the East Eurasian cohort.

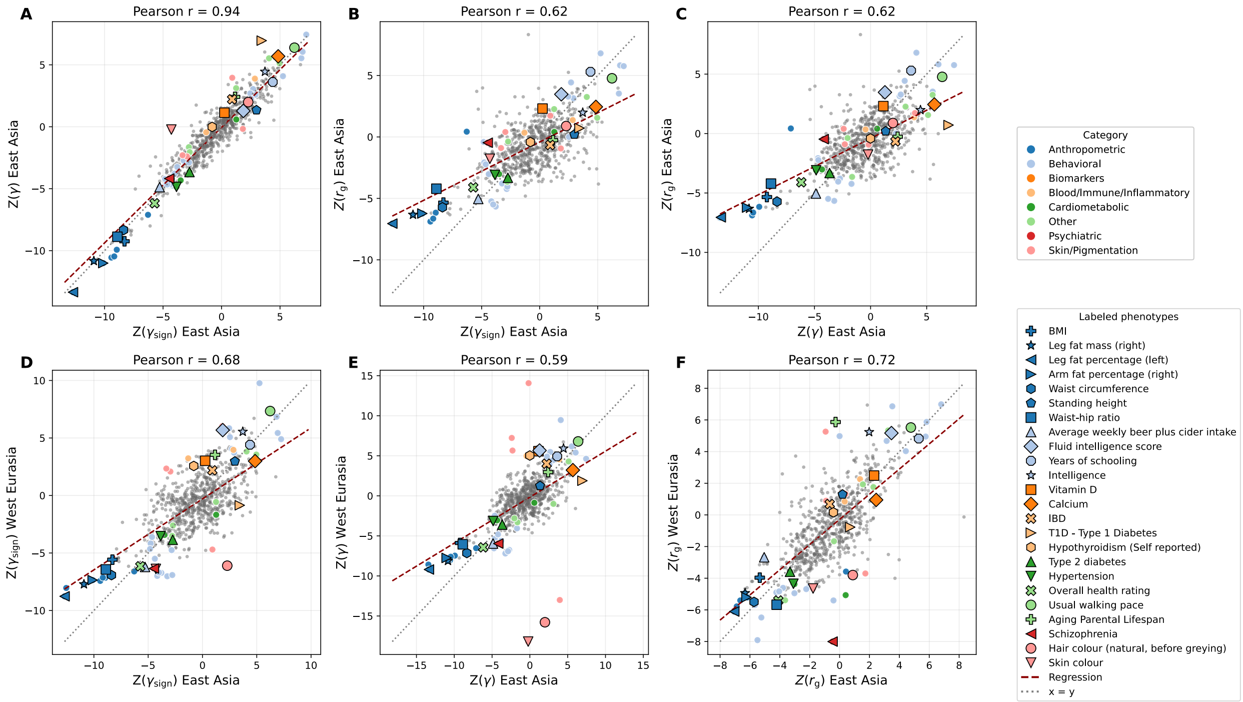


**Fig. S3.**

Consistency in polygenic tests of selection is maintained when restricting to a high East Asian ancestry cohort (**Supplementary Text**). **(A-C)** There is strong correlation between the 3 tests of selection from ref. (*1*) in the East Eurasian cohort including $\gamma$, $\gamma_{sign}$, and $r_s$. **(D-F)** There is correlation between each of the polygenic tests for selection in the East Eurasian and West Eurasian cohort for the same phenotypes.

**Table S1.**

**2472 modern and ancient individuals analyze.** Data Source 1 is 1605 modern (n=610) and ancient (n=995) individuals whose previously published sequences we reanalyze. Data Sources 2-4 are individuals represented by new data. Data Source 2 is 12 individuals with previously published in-solution enrichment data for which we report and analyze whole genome shotgun data. Data Source 3 is 38 individuals with previously published data for which we report and analyze additional in-solution enrichment data. Data Source 4 is 817 never-before-reported ancient individuals for which we report and analyze data that is anonymized except for a point estimate of the data of origin and information about broad region in East Eurasia. Available as an Excel table.

**Table S2.**

**16 newly reported shotgun ancient genomes.** Most are from individuals with previously reported in-solution enrichment data (n=12); the remainder are from individuals with data reported for the first time (n=4). Available as an Excel table.

**Table S3.**

**1093 ancient DNA libraries with newly reported capture data.** The majority (n=1050) are from 813 never-before-reported ancient individuals; the rest (n=43) are from 223 individuals for which we increase data quality. Available as an Excel table.

**Table S4.**

**Summary statistics for 696 tests of polygenic selection.** The data include: 452 European GWAS from Pan-UKBB[51], 107 curated European GWAS used for S-LDSC meta-analysis[66,67], 102 family GWAS from [68] (three GWAS for each trait), 2 GWAS from[69], 2 GWAS from[70], 30 East Asian GWAS from Biobank Japan[11], and one from[71]. Available as an Excel table.

**Table S5.**

**Summary statistics for 3 pigmentation phenotypes tested at different p-value thresholds for gamma and gamma_sign and rs.** The data are derived from 3 GWAS[66] on ~48k East Asian individuals for skin luminance (GCST90320257), red-green pigmentation (GCST90320258), and blue-yellow pigmentation (GCST90320259).

**Table S6.**

**Summary statistics for 696 tests of polygenic selection for the ancestry restricted cohort.** GWAS are the same as those in Supplementary Table 5. Available as an Excel table.

**Data S1.**

**Summary statistics for selection at 6.7 million variants.** The data are provided as a tab-delimited text file, compressed using gzip. Available at Harvard Dataverse upon final publication.

**Data S2.**

**Summary statistics for selection at 6.7 million variants for the ancestry restricted cohort.** The data are provided as a tab-delimited text file, compressed using gzip. Available at Harvard Dataverse. Available at Harvard Dataverse upon final publication

12. H.-C. Yang, P.-Y. Kwok, L.-H. Li, Y.-M. Liu, Y.-J. Jong, K.-Y. Lee, D.-W. Wang, M.-F. Tsai, J.-H. Yang, C.-H. Chen, E.-C. Yeh, C. Wei, C. S.-J. Fann, Y.-T. Huang, C.-W. Chen, Y.-J. Lee, S.-K. Chu, C. Ho, C.-S. Yang, Y. L. Lee, H.-H. Chen, M.-C. Hou, J.-F. Chiou, S.-F. Yang, C.-H. Wang, C.-Y. Huang, K.-M. Chiu, M. Chen, F.-T. Chiang, S.-L. Lee, S.-S. Chen, W.-J. Yao, C.-C. Chien, S.-Y. Lin, F.-P. Chang, H.-L. Ho, Y.-C. Yeh, W.-C. Tseng, M.-H. Lin, H.-T. Chang, L.-M. Tseng, W.-Y. Liang, P. C.-H. Chen, J.-F. Hang, S.-C. Lin, Y.-J. Chan, Y.-J. Kuo, L.-C. Wang, C.-C. Pan, Y.-C. Hsieh, Y.-M. Chen, T.-H. Hsiao, C.-H. Lin, Y.-J. Chen, I.-C. Chen, C.-L. Mao, S.-J. Chang, Y.-L. Chang, Y.-J. Liao, C.-H. Lai, W.-J. Lee, H. Tung, T.-T. Yen, H.-C. Yen, C.-M. Shih, T.-Y. Chou, T.-H. Liou, C.-Y. Chiang, Y.-G. Cherng, C.-H. Chen, C.-H. Chiu, S.-H. Tseng, E. P.-Y. Lin, Y.-J. Chen, H.-P. Chuang, T.-C. Chen, W.-T. Huang, J. Sin, I.-L. Liu, Y.-C. Chen, K.-K. Chao, Y.-M. Wu, P.-P. Yu, L.-P. Chang, K.-Y. Yen, L.-C. Chang, Y.-J. Sheen, Y.-T. Chen, K. Kan, H.-L. Tsai, Y.-K. Wang, M.-F. Hou, Y.-H. Yang, C.-H. Kuo, W.-J. Wu, J.-F. Huang, I.-W. Chong, J.-R. Tsai, C.-Y. Lin, M.-C. Yu, T.-H. Lee, M.-H. Tsai, Y.-C. Ou, P.-Y. Chen, T.-H. Hu, Y.-C. Shyu, C.-K. Cheng, Y.-J. Fang, S.-C. Hsieh, C.-H. Chen, C.-C. Chen, K.-J. Li, C.-H. Lin, H.-Y. Chiu, C.-C. Wu, C.-Y. Chen, S.-J. Chu, F.-C. Liu, F.-C. Yang, H.-A. Chang, W. Chen, S.-S. Yang, Y. Sung, T.-F. Wang, S.-Z. Lin, Y.-W. Wu, C.-S. Wu, J.-Y. Jiang, G.-C. Ma, T.-Y. Chang, J.-J. Hwang, K.-J. Kao, C.-F. Hung, T.-F. Chiu, P.-Y. Chen, K. Tsui, M.-S. Wu, S.-T. Pang, S.-A. Chen, W.-M. Chen, C. Chen, W. H.-H. Sheu, J.-Y. Wu, The Taiwan Precision Medicine Initiative provides a cohort for large-scale studies. *Nature*, 1–11 (2025).

35. M. V. Holmes, C. E. Dale, L. Zuccolo, R. J. Silverwood, Y. Guo, Z. Ye, D. Prieto-Merino, A. Dehghan, S. Trompet, A. Wong, A. Cavadino, D. Drogan, S. Padmanabhan, S. Li, A. Yesupriya, M. Leusink, J. Sundstrom, J. A. Hubacek, H. Pikhart, D. I. Swerdlow, A. G. Panayiotou, S. A. Borinskaya, C. Finan, S. Shah, K. B. Kuchenbaecker, T. Shah, J. Engmann, L. Folkersen, P. Eriksson, F. Ricceri, O. Melander, C. Sacerdote, D. M. Gamble, S. Rayaprolu, O. A. Ross, S. McLachlan, O. Vikhireva, I. Sluijs, R. A. Scott, V. Adamkova, L. Flicker, F. M. van Bockxmeer, C. Power, P. Marques-Vidal, T. Meade, M. G. Marmot, J. M. Ferro, S. Paulos-Pinheiro, S. E. Humphries, P. J. Talmud, I. M. Leach, N. Verweij, A. Linneberg, T. Skaaby, P. A. Doevendans, M. J. Cramer, P. van der Harst, O. H. Klungel, N. F. Dowling, A. F. Dominiczak, M. Kumari, A. N. Nicolaides, C. Weikert, H. Boeing, S. Ebrahim, T. R. Gaunt, J. F. Price, L. Lannfelt, A. Peasey, R. Kubinova, A. Pajak, S. Malyutina, M. I. Voevoda, A. Tamosiunas, A. H. M. der Zee, P. E. Norman, G. J. Hankey, M. M. Bergmann, A. Hofman, O. H. Franco, J. Cooper, J. Palmen, W. Spiering, P. A. de Jong, D. Kuh, R. Hardy, A. G. Uitterlinden, M. A. Ikram, I. Ford, E. Hyppönen, O. P. Almeida, N. J. Wareham, K.-T. Khaw, A. Hamsten, L. L. N. Husemoen, A. Tjønneland, J. S. Tolstrup, E. Rimm, J. W. J. Beulens, W. M. M. Verschuren, N. C. Onland-Moret, M. H. Hofker, S. G. Wannamethee, P. H. Whincup, R. Morris, A. M. Vicente, H. Watkins, M. Farrall, J. W. Jukema, J. Meschia, L. A. Cupples, S. J. Sharp, M. Fornage, C. Kooperberg,

A. Z. LaCroix, J. Y. Dai, M. B. Lanktree, D. S. Siscovick, E. Jorgenson, B. Spring, J. Coresh, Y. R. Li, S. G. Buxbaum, P. J. Schreiner, R. C. Ellison, M. Y. Tsai, S. R. Patel, S. Redline, A. D. Johnson, R. C. Hoogeveen, H. Hakonarson, J. I. Rotter, E. Boerwinkle, P. I. W. de Bakker, M. Kivimaki, F. W. Asselbergs, N. Sattar, D. A. Lawlor, J. Whittaker, G. D. Smith, K. Mukamal, B. M. Psaty, J. G. Wilson, L. A. Lange, A. Hamidovic, A. D. Hingorani, B. G. Nordestgaard, M. Bobak, D. A. Leon, C. Langenberg, T. M. Palmer, A. P. Reiner, B. J. Keating, F. Dudbridge, J. P. Casas, Association between alcohol and cardiovascular disease: Mendelian randomisation analysis based on individual participant data. *BMJ* **349**, g4164 (2014).

41. K. H. Darwin, S. A. Stanley, The aldehyde hypothesis: metabolic intermediates as antimicrobial effectors. *Open Biol.* **12**, 220010.

42. A. Auton, G. R. Abecasis, D. M. Altshuler, R. M. Durbin, G. R. Abecasis, D. R. Bentley, A. Chakravarti, A. G. Clark, P. Donnelly, E. E. Eichler, P. Flicek, S. B. Gabriel, R. A. Gibbs, E. D. Green, M. E. Hurles, B. M. Knoppers, J. O. Korbel, E. S. Lander, C. Lee, H. Lehrach, E. R. Mardis, G. T. Marth, G. A. McVean, D. A. Nickerson, J. P. Schmidt, S. T. Sherry, J. Wang, R. K. Wilson, R. A. Gibbs, E. Boerwinkle, H. Doddapaneni, Y. Han, V. Korchina, C. Kovar, S. Lee, D. Muzny, J. G. Reid, Y. Zhu, J. Wang, Y. Chang, Q. Feng, X. Fang, X. Guo, M. Jian, H. Jiang, X. Jin, T. Lan, G. Li, J. Li, Y. Li, S. Liu, X. Liu, Y. Lu, X. Ma, M. Tang, B. Wang, G. Wang, H. Wu, R. Wu, X. Xu, Y. Yin, D. Zhang, W. Zhang, J. Zhao, M. Zhao, X. Zheng, E. S. Lander, D. M. Altshuler, S. B. Gabriel, N. Gupta, N. Gharani, L. H. Toji, N. P. Gerry, A. M. Resch, P. Flicek, J. Barker, L. Clarke, L. Gil, S. E. Hunt, G. Kelman, E. Kulesha, R. Leinonen, W. M. McLaren, R. Radhakrishnan, A. Roa, D. Smirnov, R. E. Smith, I. Streeter, A. Thormann, I. Toneva, B. Vaughan, X. Zheng-Bradley, D. R. Bentley, R. Grocock, S. Humphray, T. James, Z. Kingsbury, H. Lehrach, R. Sudbrak, M. W. Albrecht, V. S. Amstislavskiy, T. A. Borodina, M. Lienhard, F. Mertes, M. Sultan,

B. Timmermann, M.-L. Yaspo, E. R. Mardis, R. K. Wilson, L. Fulton, R. Fulton, S. T. Sherry, V. Ananiev, Z. Belaia, D. Beloslyudtsev, N. Bouk, C. Chen, D. Church, R. Cohen, C. Cook, J. Garner, T. Hefferon, M. Kimelman, C. Liu, J. Lopez, P. Meric, C. O'Sullivan, Y. Ostapchuk, L. Phan, S. Ponomarov, V. Schneider, E. Shekhtman, K. Sirotkin, D. Slotta, H. Zhang, G. A. McVean, R. M. Durbin, S. Balasubramaniam, J. Burton, P. Danecek, T. M. Keane, A. Kolb-Kokocinski, S. McCarthy, J. Stalker, M. Quail, J. P. Schmidt, C. J. Davies, J. Gollub, T. Webster, B. Wong, Y. Zhan, A. Auton, C. L. Campbell, Y. Kong, A. Marcketta, R. A. Gibbs, F. Yu, L. Antunes, M. Bainbridge, D. Muzny, A. Sabo, Z. Huang, J. Wang, L. J. M. Coin, L. Fang, X. Guo, X. Jin, G. Li, Q. Li, Y. Li, Z. Li, H. Lin, B. Liu, R. Luo, H. Shao, Y. Xie, C. Ye, C. Yu, F. Zhang, H. Zheng, H. Zhu, C. Alkan, E. Dal, F. Kahveci, G. T. Marth, E. P. Garrison, D. Kural, W.-P. Lee, W. Fung Leong, M. Stromberg, A. N. Ward, J. Wu, M. Zhang, M. J. Daly, M. A. DePristo, R. E. Handsaker, D. M. Altshuler, E. Banks, G. Bhatia, G. del Angel, S. B. Gabriel, G. Genovese, N. Gupta, H. Li, S. Kashin, E. S. Lander, S. A. McCarroll, J. C. Nemesh, R. E. Poplin, S. C. Yoon, J. Lihm, V. Makarov, A. G. Clark, S. Gottipati, A. Keinan, J. L. Rodriguez-Flores, J. O. Korbel, T. Rausch, M. H. Fritz, A. M. Stütz, P. Flicek, K. Beal, L. Clarke, A. Datta, J. Herrero, W. M. McLaren, G. R. S. Ritchie, R. E. Smith, D. Zerbino, X. Zheng-Bradley, P. C. Sabeti, I. Shlyakhter, S. F. Schaffner, J. Vitti, D. N. Cooper, E. V. Ball, P. D. Stenson, D. R. Bentley, B. Barnes, M. Bauer, R. Keira Cheetham, A. Cox, M. Eberle, S. Humphray, S. Kahn, L. Murray, J. Peden, R. Shaw, E. E. Kenny, M. A. Batzer, M. K. Konkel, J. A. Walker, D. G. MacArthur, M. Lek, R. Sudbrak, V. S. Amstislavskiy, R. Herwig, E. R. Mardis, L. Ding, D. C. Koboldt, D. Larson, K. Ye, S. Gravel, The 1000 Genomes Project Consortium, Corresponding authors, Steering committee, Production group, Baylor College of Medicine, BGI-Shenzhen, Broad Institute of MIT and Harvard, Coriell Institute for Medical Research, E. B. I. European Molecular Biology Laboratory, Illumina, Max Planck Institute for Molecular Genetics, McDonnell Genome Institute at Washington University, US National Institutes of Health, University of Oxford, Wellcome Trust Sanger Institute, Analysis group, Affymetrix, Albert Einstein College of Medicine, Bilkent University, Boston College, Cold Spring Harbor Laboratory, Cornell University, European Molecular Biology Laboratory, Harvard University, Human Gene Mutation Database, Icahn School of Medicine at Mount Sinai, Louisiana State University, Massachusetts General Hospital, McGill University, N. National Eye Institute, A global reference for human genetic variation. *Nature* **526**, 68–74 (2015).

56. M. Byrska-Bishop, U. S. Evani, X. Zhao, A. O. Basile, H. J. Abel, A. A. Regier, A. Corvelo, W. E. Clarke, R. Musunuri, K. Nagulapalli, S. Fairley, A. Runnels, L. Winterkorn,

E. Lowy, Human Genome Structural Variation Consortium,  null Paul Flicek, S. Germer, H. Brand, I. M. Hall, M. E. Talkowski, G. Narzisi, M. C. Zody, High-coverage whole-genome sequencing of the expanded 1000 Genomes Project cohort including 602 trios. *Cell* **185**, 3426-3440.e19 (2022).

57. S. Rubinacci, D. M. Ribeiro, R. J. Hofmeister, O. Delaneau, Efficient phasing and imputation of low-coverage sequencing data using large reference panels. *Nat. Genet.* **53**, 120–126 (2021).

58. P. Danecek, J. K. Bonfield, J. Liddle, J. Marshall, V. Ohan, M. O. Pollard, A. Whitwham, T. Keane, S. A. McCarthy, R. M. Davies, H. Li, Twelve years of SAMtools and BCFtools. *GigaScience* **10**, giab008 (2021).

59. H. Zhao, Z. Sun, J. Wang, H. Huang, J.-P. Kocher, L. Wang, CrossMap: a versatile tool for coordinate conversion between genome assemblies. *Bioinformatics* **30**, 1006–1007 (2014).

60. S. Chen, L. C. Francioli, J. K. Goodrich, R. L. Collins, M. Kanai, Q. Wang, J. Alföldi, N. A. Watts, C. Vittal, L. D. Gauthier, T. Poterba, M. W. Wilson, Y. Tarasova, W. Phu, R. Grant, M. T. Yohannes, Z. Koenig, Y. Farjoun, E. Banks, S. Donnelly, S. Gabriel, N. Gupta, S. Ferriera, C. Tolonen, S. Novod, L. Bergelson, D. Roazen, V. Ruano-Rubio, M. Covarrubias, C. Llanwarne, N. Petrillo, G. Wade, T. Jeandet, R. Munshi, K. Tibbetts, Genome Aggregation Database Consortium, A. O'Donnell-Luria, M. Solomonson, C. Seed, A. R. Martin, M. E. Talkowski, H. L. Rehm, M. J. Daly, G. Tiao, B. M. Neale, D. G. MacArthur, K. J. Karczewski, A genomic mutational constraint map using variation in 76,156 human genomes. *Nature* **625**, 92–100 (2024).

69. M. Larena, F. Sanchez-Quinto, P. Sjödin, J. McKenna, C. Ebeo, R. Reyes, O. Casel, J.-Y. Huang, K. P. Hagada, D. Guilay, J. Reyes, F. P. Allian, V. Mori, L. S. Azarcon, A. Manera, C. Terando, L. Jamero, G. Sireg, R. Manginsay-Tremedal, M. S. Labos, R. D. Vilar, A. Latiph, R. L. Saway, E. Marte, P. Magbanua, A. Morales, I. Java, R. Reveche, B. Barrios, E. Burton, J. C. Salon, Ma. J. T. Kels, A. Albano, R. B. Cruz-Angeles, E. Molanida, L. Granehäll, M. Vicente, H. Edlund, J.-H. Loo, J. Trejaut, S. Y. W. Ho, L. Reid, H. Malmström, C. Schlebusch, K. Lambeck, P. Endicott, M. Jakobsson, Multiple migrations to the Philippines during the last 50,000 years. *Proc. Natl. Acad. Sci.* **118**, e2026132118 (2021).

70. A. S. Hinrichs, D. Karolchik, R. Baertsch, G. P. Barber, G. Bejerano, H. Clawson, M. Diekhans, T. S. Furey, R. A. Harte, F. Hsu, J. Hillman-Jackson, R. M. Kuhn, J. S. Pedersen, A. Pohl, B. J. Raney, K. R. Rosenbloom, A. Siepel, K. E. Smith, C. W. Sugnet, A. Sultan-Qurraie, D. J. Thomas, H. Trumbower, R. J. Weber, M. Weirauch, A. S. Zweig, D. Haussler, W. J. Kent, The UCSC Genome Browser Database: update 2006. *Nucleic Acids Res.* **34**, D590-598 (2006).
